# Thrombopoietin signaling to chromatin elicits rapid and pervasive epigenome remodeling within poised chromatin architectures

**DOI:** 10.1101/163113

**Authors:** Federico Comoglio, Hyun Jung Park, Stefan Schoenfelder, Iros Barozzi, Daniel Bode, Peter Fraser, Anthony R. Green

**Author notes:** These authors contributed equally to this work (Federico Comoglio, Hyun Jung Park). Correspondence (F.C.); (A.R.G.).

## Abstract

Thrombopoietin (TPO) is a critical cytokine regulating hematopoietic stem cell maintenance and differentiation into the megakaryocytic lineage. However, the transcriptional and chromatin dynamics elicited by TPO signaling are poorly understood. Here, we study the immediate early transcriptional and cis-regulatory responses to TPO in hematopoietic stem/progenitor cells (HSPCs) and use this paradigm of cytokine signaling to chromatin to dissect the relation between cis-regulatory activity and chromatin architecture. We show that TPO profoundly alters the transcriptome of HSPCs, with key hematopoietic regulators being transcriptionally repressed within 30 minutes of TPO. By examining cis-regulatory dynamics and chromatin architectures, we demonstrate that these changes are accompanied by rapid and extensive epigenome remodeling of cis-regulatory landscapes that is spatially coordinated within topologically associating domains (TADs). Moreover, TPO-responsive enhancers are spatially clustered and engage in preferential homotypic intra‐ and inter-TAD interactions that are largely refractory to TPO signaling. By further examining the link between cis-regulatory dynamics and chromatin looping, we show that rapid modulation of cis-regulatory activity is largely independent of chromatin looping dynamics. Finally, we show that, although activated and repressed cis-regulatory elements share remarkably similar DNA sequence compositions, transcription factor binding patterns accurately predict rapid cis-regulatory responses to TPO.

## Introduction

Hematopoiesis - the formation of blood cellular components - is an exquisitely characterized process in which the signaling consequences of extracellular stimuli, such as cytokines and growth factors, must be coherently integrated with chromatin structures to regulate cell-type-specific transcriptional programs (Rieger and Schroeder 2012). Specification of blood cell types is thought to proceed in a hierarchical fashion. At the apex of the hematopoietic hierarchy are multipotent hematopoietic stem cells (HSCs), which are able both to self-renew and to generate all blood cell lineages through differentiation into increasingly mature progenitor cells (Eaves 2015).

Lineage choice and commitment throughout hematopoiesis entails induction of lineage-specific gene regulatory networks and repression of lineage-inappropriate genes (Rieger and Schroeder 2012). These transcriptional programs are the result of coordinated waves of activation and de-commissioning of unique constellations of cis-regulatory elements such as promoters and enhancers (Spitz and Furlong 2012). Cell-type-specific access to cis-regulatory information reflects the combinatorial activity of transcription factors (TFs), which recruit chromatin modifiers to establish active or repressive chromatin environments at regulatory elements within a cis-regulatory repertoire (Reiter et al. 2017).

Recent epigenomic advances have made it possible to map these repertoires genome-wide and across a wide range of cell types and organisms (Shlyueva et al. 2014; The ENCODE Project Consortium 2012). Putative enhancers can be defined as DNase I hypersensitive sites (DHSs) marked by monomethylated lysine 4 on histone H3 (H3K4me1) and their activity can be inferred based on the concomitant presence of histone H3 acetylation at lysine 27 (H3K27ac) often accompanied by detectable production of enhancer RNAs (eRNAs) (Heintzman et al. 2007, 2009; Creyghton et al. 2010; Calo and Wysocka 2013; Li et al. 2016). However, the targets of enhancers cannot be accurately predicted solely based on genomic proximity since transcriptional regulation occurs within the three-dimensional nuclear space (Bonev and Cavalli 2016; Dekker and Mirny 2016). Enhancers can exert their regulatory function on the expression of distally located target genes through three-dimensional chromatin looping that results in the juxtaposition of enhancer(s) and target promoter(s) (Bouwman and de Laat 2015; Bonev and Cavalli 2016). Experimental tethering of an enhancer to its target gene promoter has been shown to result in transcriptional activation of the *Hbb* locus, demonstrating that enhancer-promoter contacts can induce gene activation, even in the absence of a key transcriptional activator (Deng et al. 2012).

Chromosome conformation capture (3C) based methods, particularly high-throughput 3C (Hi-C), have enabled targeted or genome-wide mapping of chromatin architectures (Denker and De Laat 2016). These technologies provided critical insights into key structural and functional components of three-dimensional chromatin organization such as i) A/B compartments (Lieberman-aiden et al. 2009), also referred to as compartment domains (Rao et al. 2017), which are closely associated with open and closed chromatin domains, respectively; ii) topologically associating domains (TADs) (Dixon et al. 2012; Nora et al. 2012; Sexton et al. 2012), also referred to as contact domains (Rao et al. 2017), chromosomal units that spatially constrain cis-regulatory interactions; iii) CTCF loops, also referred to as insulated neighborhoods (Hnisz et al. 2016) or loop domains (Rao et al. 2017). Interestingly, while these studies suggested a hierarchical domain organization, recent studies based on acute depletion of CTCF or cohesin, or inactivation of the cohesin-loading factor NIPBL, demonstrated that A/B compartments and TADs are not hierarchically organized but represent independent structural (and possibly functional) units of 3D genome organization (Nora et al. 2017; Rao et al. 2017; Wutz et al. 2017; Schwarzer et al. 2017).

Enhancer-promoter communication preferentially occurs within TADs (Zhan et al. 2017). However, the high complexity of Hi-C libraries makes it impractical to systematically map cis-regulatory interactions with high resolution and coverage. To overcome this limitation, we recently developed Promoter Capture Hi-C (PCHi-C), a sequence capture approach that selectively enriches Hi-C libraries for interactions involving more than 22,000 annotated mouse promoters, thus allowing global mapping of promoter interactions at restriction fragment level resolution (Schoenfelder et al. 2015).

Collectively, studies in different systems across multiple species using 3C-based methods and derivatives identified two types of enhancer-promoter interactions: loops formed de novo and pre-existing loops (Melo et al. 2013; Eijkelenboom et al. 2014; Freire-Pritchett et al. 2017; Montavon et al. 2011; Ghavi-Helm et al. 2014; Apostolou et al. 2013; Cruz-Molina et al. 2017). While de novo (also called instructive) interactions appear concomitant with changes in the target gene activity, pre-existing (also called permissive) interactions are formed prior to gene activation and are thought to facilitate timely transcriptional induction (de Laat and Duboule 2013; Bouwman and de Laat 2015). However, the relation between signal-dependent modulation of cis-regulatory activity and chromatin looping remains poorly understood for several reasons. First, poised chromatin architectures have been primarily studied in relation to gene activation and little is known about signaling-dependent modulation of gene repression. Secondly, studies analyzing this relation on developmental timescales lack the temporal resolution necessary to capture the early chromatin changes induced by signaling events. Thirdly, physical interactions other than promoter-enhancer loops, such as enhancer-enhancer interactions, received comparatively little attention, albeit they likely represent an important layer of gene regulation in the three-dimensional nuclear space (Ing-Simmons et al. 2015).

The cytokine thrombopoietin (TPO) is a critical regulator of megakaryopoiesis. TPO regulates megakaryocyte and platelet production by activating its receptor, MPL (de Sauvage et al. 1994; Lok et al. 1994; Bartley et al. 1994), inducing multiple signaling pathways including Janus kinase (JAK)/signal transducer and activator of transcription (STAT), mitogen-activated protein kinase and phosphatidylinositol-3-kinase (De Graaf and Metcalf 2011). The physiological role of TPO has been extensively studied in mouse knockout (KO) models. TPO KO mice exhibit severe thrombocytopenia with 90% reduction in platelet counts, indicating that TPO is the major physiological regulator of megakaryocyte and platelet production in vivo (de Sauvage et al. 1996). Moreover, *Tpo* and *Mpl* KO mouse models revealed that TPO signaling is also vital for haematopoietic stem cell maintenance and self-renewal (Kimura et al. 1998; Solar et al. 1998; Buza-vidas et al. 2006; Yoshihara et al. 2007; Qian et al. 2007). However, these studies largely focused on TPO signaling at the level of cytoplasmic signaling pathways and/or cellular behavior. By contrast, the transcriptional consequences and cis-regulatory dynamics of TPO signaling to chromatin remain largely unknown.

## Results

### The immediate early transcriptional response to TPO signaling

To capture the immediate early transcriptional response to TPO, we performed subcellular RNA-seq (Bhatt et al. 2012) before and after 30 minutes TPO stimulation of HPC-7 cells, a cytokine-dependent and karyotypically normal multipotent hematopoietic stem/progenitor cell line (Pinto Do Ó et al. 1998). HPC-7 cells self-renew in the presence of stem cell factor (SCF) and undergo megakaryocytic differentiation following TPO stimulation.

We isolated and profiled biochemically fractionated chromatin-associated, nucleoplasmic, and cytoplasmic transcripts in two biological replicates. Principal component analysis and hierarchical clustering indicate high reproducibility between replicates, with the subcellular compartment explaining most of the observed variation in gene expression (PC1, 72%) (Supplemental Fig. S1A-B). As expected, this is largely driven by the high intronic content of chromatin-associated RNA-seq libraries, which are enriched for unprocessed and incompletely spliced transcripts (Supplemental Fig. S1C). In addition, analysis of small nuclear and small cytoplasmic RNA gene loci revealed compartment-specific transcript localization, confirming the high enrichment of our RNA fractions (Supplemental Fig. S1D).

We then analyzed the immediate early transcriptional consequences of TPO signaling at the chromatin level. We found that TPO transcriptionally upregulated 1,325 genes and repressed 639 genes (Fig.1A). Importantly, significant changes in chromatin-associated RNA levels within 30’ of TPO stimulation only moderately overlapped changes detected in the cytoplasmic fraction (58% and 37% for up‐ and down-regulated genes, respectively) (Supplemental Fig. S1E) and exhibited significantly higher fold changes (Supplemental Fig. S1F). This suggests that RNA stability and post-transcriptional regulation contribute substantially to cytoplasmic gene expression changes even in response to transient signaling, underscoring the importance of profiling chromatin-associated or nascent transcripts when studying primary transcriptional responses. Unless otherwise stated the remainder of our expression analyses focused on chromatin-associated transcripts.

**Figure 1.**
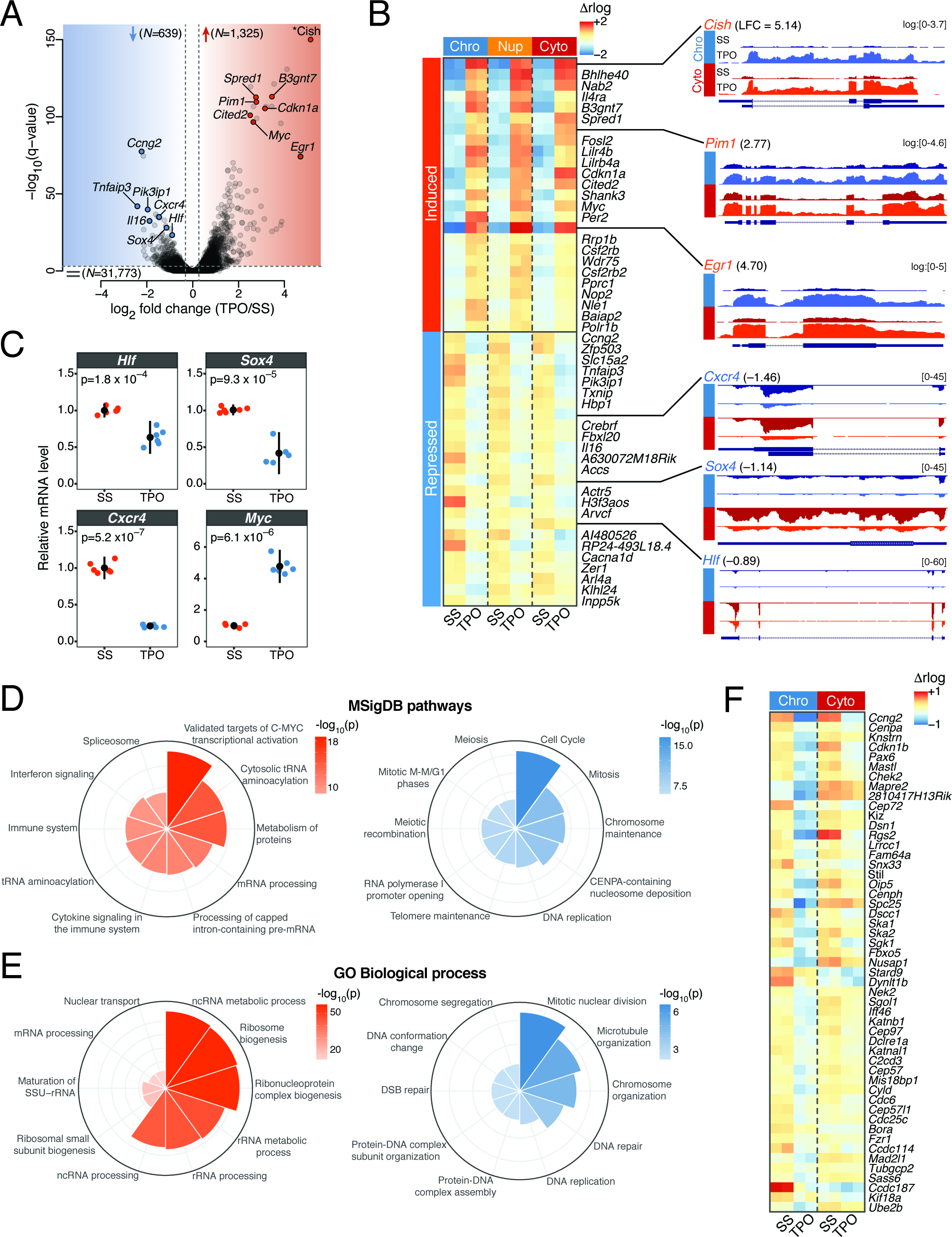
The immediate early transcriptional response to TPO signaling. **(A)** Volcano plot of gene transcription changes induced by 30’ stimulation of HPC-7 cells with TPO relative to serum-starved (SS) control cells, based on chromatin-associated RNA expression. Red and blue shaded regions enclose transcriptionally up‐ and down-regulated genes (q-value < 10^−3^), respectively. The total number of genes within each category is indicated. Representative hits are labeled. The asterisk denotes the most strongly induced gene (*Cish*, q = 1.8 x 10^−241^), which was repositioned within the plot area. **(B)** (left) Chromatin-associated (Chro), nucleoplasmic (Nup) and cytoplasmic (Cyto) RNA expression heatmap for the top 25 induced and top 25 repressed genes ranked by q-value. Regularized log2 (rlog) expression values are row-mean subtracted. (right) Representative tracks of differentially transcribed genes. Where indicated, RNA-seq coverage was log transformed with a pseudocount of 1. LFC, log2 fold change. **(C)** mRNA expression levels of the indicated genes in TPO-treated (30’) primary CD41^+^LSK cells, relative to serum-starved (SS) control cells, measured by quantitative RT-PCR. Error bars are mean±SD (n=6) from two mice. P-values are from a two-sided Welch’s t-test. **(D)** Top 10 significantly enriched Molecular Signature Database (MSigDB) pathways for transcriptionally up-(left) and down-regulated (right) genes, ranked by binomial p-value. **(E)** Same as (D), for Gene Ontology (GO) biological process terms. **(F)** Chromatin-associated (Chro) and cytoplasmic (Cyto) RNA expression heatmap of mitotic genes. Regularized log2 (rlog) expression values are row-mean subtracted.

Interestingly, the immediate early transcriptional consequences of TPO signaling to chromatin were highly divergent. On the one hand, genes most highly transcriptionally upregulated by TPO included several targets of the canonical JAK/STAT signaling pathway (e.g. *Cish*, *Pim1*, *Cited2* and *Egr1*) and the transcription factor MYC (Fig. 1A-B), suggesting broad housekeeping and survival functions. On the other hand, TPO led to the rapid transcriptional repression of key hematopoietic regulators such as *Hlf*, *Sox4* and *Cxcr4* (Fig. 1B). Repression of these loci resulted in strongly decreased cytoplasmic RNA levels within 30 minutes of TPO, indicating that transcripts encoding these key regulators are subjected to rapid turnover. To validate our results in primary cells, we confirmed the rapid TPO-induced downregulation of *Hlf*, *Sox4* and *Cxcr4*, as well as upregulation of *Myc* by RT-qPCR in CD41^+^Lin^-^ Sca1^+^c-Kit^+^ (CD41^+^LSK) bone marrow cells (Nishikii et al. 2015) (Fig. 1C), indicating that TPO elicits a common transcriptional program in megakaryocytic-biased hematopoietic progenitors.

To gain further insights into the nature of the transcriptional programs regulated by TPO, we subjected differentially transcribed genes to a Molecular Signatures Database (MSigDB) and Gene Ontology (GO) enrichment analyses. Transcriptionally upregulated events were strongly enriched for housekeeping genes involved in RNA and protein metabolism whose expression is largely driven by a MYC transcriptional program, and for genes that respond to cytokine signaling in the immune system (Fig. 1D-E). However, megakaryocytic-affiliated genes were not induced within 30 minutes of TPO (Supplemental Fig. S1A-D and S2), consistent with a slower induction kinetic (Park et al. 2015). In contrast, genes involved in mitosis, chromosome maintenance and DNA repair were rapidly repressed by TPO (Fig. 1D-F). A distinctive feature of megakaryopoiesis is endomitosis, DNA replication in the absence of cell division that results from an incomplete M phase due to a failure in late cytokinesis (Bluteau et al. 2009). The rapid transcriptional repression of genes involved in mitotic nuclear division and microtubule organization (Fig. 1F) suggests that a cell cycle switch to endomitosis might occur very rapidly at the transcriptional level. Alternatively, it might represent a more general phenomenon of cell differentiation (Ruijtenberg and Heuvel 2016).

### TPO signaling elicits rapid and extensive epigenome remodeling of cis-regulatory landscapes

Next, we set out to investigate the chromatin dynamics underlying the immediate early response to TPO by surveying the activity of cis-regulatory elements before and after 30 minutes TPO stimulation of HPC-7 cells. To this end, we profiled H3K27ac genome-wide by chromatin immunoprecipitation sequencing (ChIP-seq) in two biological replicates and used a sliding window-based approach to detect genomic regions exhibiting significantly altered H3K27ac levels at 1% false discovery rate (FDR) (Fig. 2A, see Methods) (Lun and Smyth 2015).

**Figure 2.**
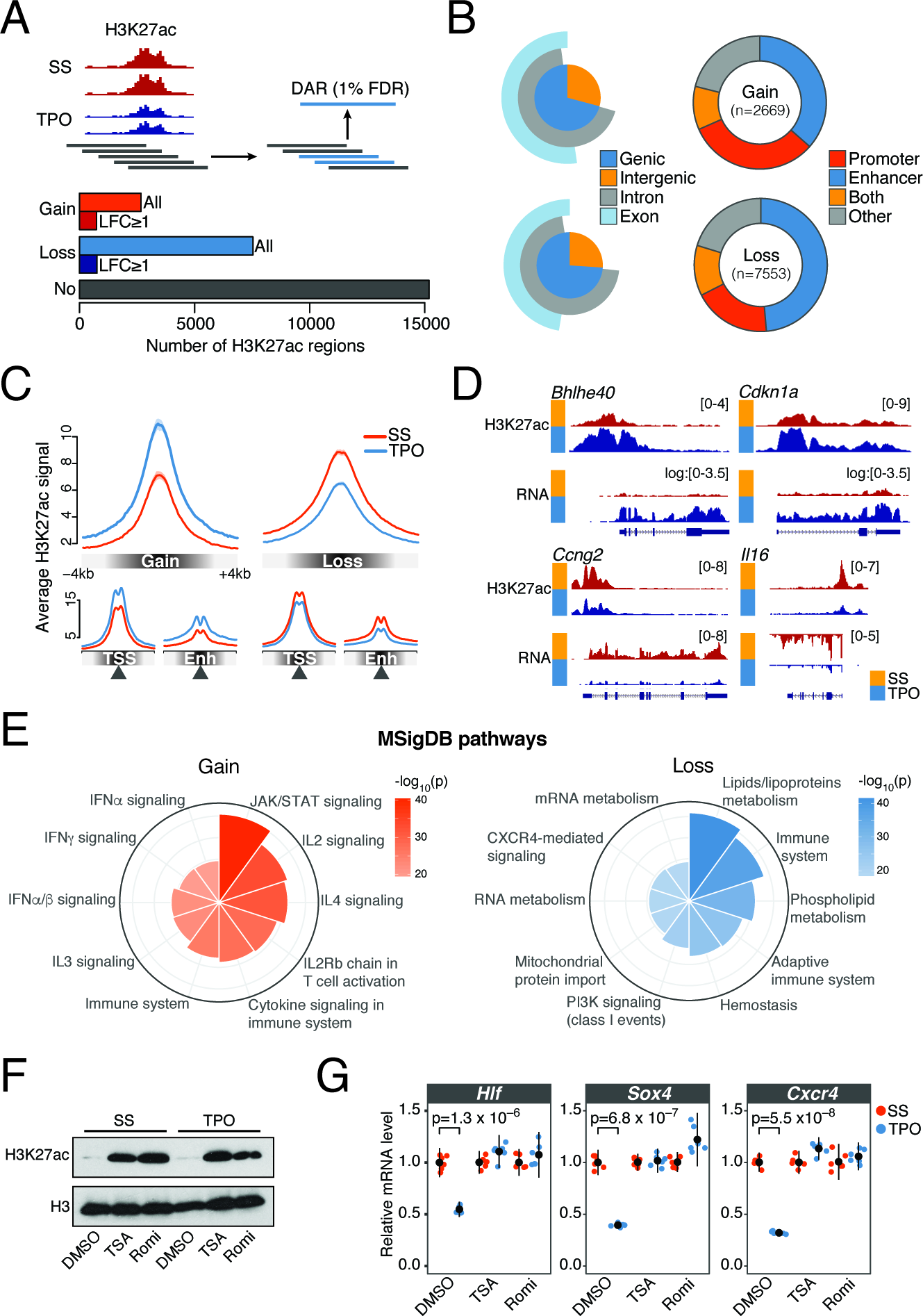
TPO signaling elicits rapid and extensive epigenome remodeling at cis-regulatory elements. **(A)** (top) Schematic representation of the tiling-window approach used to identify differentially acetylated regions (DARs) from H3K27ac ChIP-seq profiles. Significant overlapping windows (blue rectangles) were merged to call DARs at 1% false discovery rate (FDR). (bottom) Total number of activated (Gain) and repressed (Loss) DARs. H3K27ac regions not significantly altered by 30’ TPO (No) are also shown. LFC, log2 fold change. **(B)** (left) Annotation of DARs with respect to genomic compartments. (right) Annotation of DARs with respect to cis-regulatory elements inferred from DNase-seq profiles in serum-starved HPC-7 cells (see Methods). **(C)** Average normalized H3K27ac signal within ±4 kb of DAR summits (top), transcription start sites (TSSs) or enhancer summits (Enh). **(D)** Representative tracks of DARs. Where indicated, chromatin-associated RNA-seq coverage was log transformed with a pseudocount of 1. **(E)** Genomic regions enrichment of annotations tool (GREAT) analysis of DARs. Top 10 significantly enriched Molecular Signature Database (MSigDB) pathways for gene associated with DARs, ranked by binomial p-value. **(F)** Western blot analysis of H3K27ac and total H3 levels in HPC-7 cells pre-treated for 30 minutes with DMSO, Trichostatin A (TSA) or Romidepsin (Romi), before and after 30 minutes TPO stimulation. **(G)** Corresponding mRNA expression levels of the indicated genes relative to serum-starved (SS) control cells, measured by quantitative RT-PCR. Error bars are mean±SD (n=6) from two biological replicates. p-values are from a two-sided Welch’s *t*-test.

Our analysis identified 10,222 differentially acetylated regions (DARs) exhibiting significantly increased (Gain, n = 2,669) or decreased (Loss, n = 7,553) H3K27ac levels within 30’ of TPO treatment (Fig. 2A). This number represents a sizeable fraction (40%) of genomic regions marked by H3K27ac in basal condition, demonstrating widespread (albeit not global) changes of cis-regulatory activity in response to transient cytokine signaling (Fig. 2A). Activated and repressed DARs showed overlapping size distributions with a median length of 3.4kb (Supplemental Fig. S3A). However, the two sets differed in the magnitude of their response to TPO, with activated DARs exhibiting significantly stronger changes in H3K27ac levels (Fig. 2A and Supplemental Fig. S3B). We ruled out the possibility that these broad acetylation changes could stem from technical biases, as H3K27ac promoter signals at the 1,000 most highly expressed genes unaffected by TPO were highly correlated (r=0.94-0.99) and exhibited no systematic difference between replicates and conditions (Supplemental Fig. S3C). In addition, changes in the abundance of enhancer RNA (eRNA) transcripts at differentially acetylated intergenic enhancers correlated with changes in H3K27ac levels, further validating our DAR calls (Supplemental Fig. S3D). Moreover, to rule out the possibility that differential acetylation events reflect global changes in H3 occupancy, we measured H3K27ac and total H3 levels at a set of differentially acetylated loci by ChIP-qPCR (Supplemental Fig. S3E). This analysis indicates that H3K27ac changes at DARs represent bona fide acetylation dynamics.

We next mapped DARs onto genomic compartments. Less than 30% of DARs were intergenic, with the majority of acetylation changes overlapping annotated genes (Fig. 2B). To more precisely localize cis-regulatory elements within DARs, we mapped active promoters and enhancers at high resolution using DNase-seq profiles in self-renewing HPC-7 cells (see Methods) (Wilson et al. 2016). The vast majority (80%) of DARs encompassed elements (DHSs) from these two regulatory classes (Fig. 2B-D).

To investigate the functional significance of DARs, we performed a Genomic Regions Enrichment of Annotations Tool (GREAT) analysis by linking DARs to their putative target genes (McLean et al. 2010). As expected, activated DARs were strongly linked with genes involved in cytokine and JAK/STAT signaling (Fig. 2E). In contrast, repressed DARs were significantly associated to genes involved in lipid metabolism and in the regulation of the immune system, particularly of adaptive immunity (Fig. 2E), indicating that TPO signaling represses the cis-regulatory elements of lymphoid-affiliated genes.

Finally, to test whether deacetylation of cis-regulatory elements is required for transcriptional repression of key hematopoietic regulators (Fig. 1B-C), we pre-treated HPC-7 cells with DMSO or the HDAC inhibitors Trichostatin A (TSA) or Romidepsin (Romi) for 30 minutes (Fig. 2F) and measured gene expression changes induced by 30 minutes TPO stimulation by RT-qPCR. We found that whereas *Hlf*, *Sox4* and *Cxcr4* were significantly downregulated within 30 minutes of TPO treatment in DMSO-treated cells, TSA or Romi fully abrogated this response (Fig. 2G).

Together, these results demonstrate that TPO signaling elicits rapid and extensive epigenome remodeling at cis-regulatory elements and that histone deacetylation is necessary for transcriptional repression of key hematopoietic regulators.

### Cis-regulatory dynamics induced by TPO signaling are spatially coordinated within topological domains

We noted that homotypic DARs tended to strongly cluster along the chromatin fiber, suggesting spatial coordination within chromatin domains (Fig. 3A). Thus, we set out to systematically investigate how the rapid modulation of cis-regulatory activity induced by TPO is spatially coordinated within the nucleus. To this end, we generated Hi-C libraries from HPC-7 cells before and after 30 minutes TPO stimulation in two biological replicates and identified valid di-tags using the Hi-C Pro pipeline (Servant et al. 2015). We obtained a of total of 644 and 675 million valid ditags in basal and TPO-treated conditions, respectively. Of these, 565 (87.7%) and 593 (87.9%) million valid di-tags, respectively, were intrachromosomal paired-end reads.

**Figure 3.**
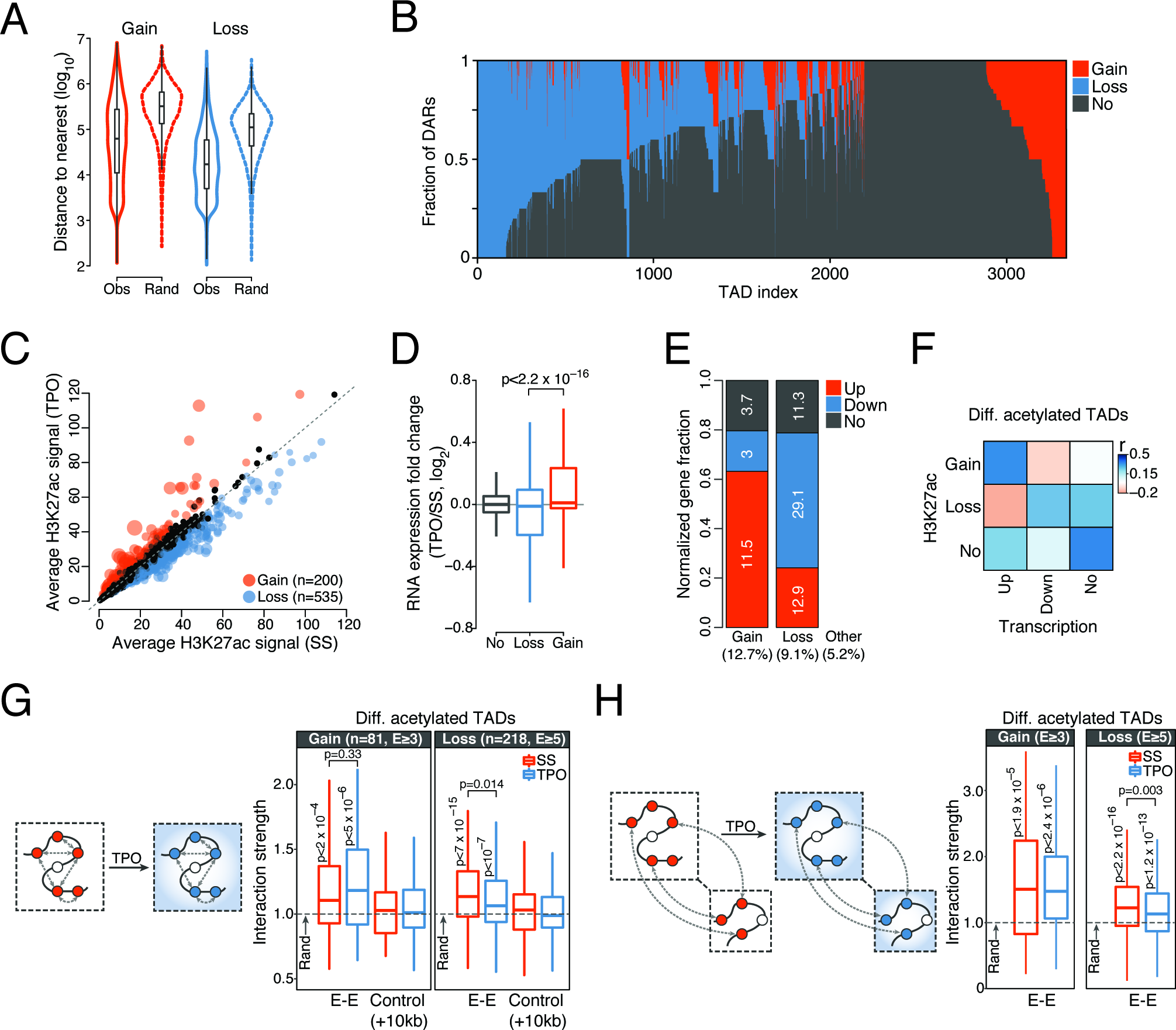
is-regulatory responses to TPO are spatially coordinated within the nucleus. **(A)** Distribution of observed (Obs) and expected (Rand, see Methods) genomic distances between nearest homotypic DARs. **(B)** Distribution of DARs within TADs detected in serum-starved HPC-7 cells and containing at least one H3K27ac region. Each TAD corresponds to a vertical line. **(C)** Average normalized H3K27ac signals per kb of TAD. Significantly differentially acetylated TADs are highlighted. Point sizes are proportional to ‐log_10_(q-value). **(D)** Distribution of chromatin-associated RNA expression fold changes of genes localized within differentially acetylated TADs. p-value is from a Wilcoxon rank sum test. **(E)** Relative fraction of differentially transcribed genes localized within differentially acetylated TADs, normalized by total number of genes within each category. The percentage of genes in each category is indicated inside the bar plot, whereas the percentage of differentially transcribed genes falling within each TAD class is indicated at the bottom. **(F)** Spearman’s rank correlation matrix between frequency of DAR and frequency of differentially transcribed genes within all TADs detected in serum-starved HPC-7 cells and containing at least one H3K27ac region (top) or within differentially acetylated TADs (bottom). **(G)** Structured interaction matrix analysis (SIMA) of enhancer-enhancer interactions for differentially acetylated enhancers within differentially acetylated TADs. The interaction strength reflects the enrichment of Hi-C interactions relative to randomly sampled genomic regions. Interaction strength distributions for matched controls (see Methods) are also shown. p-values are from a Wilcoxon signed rank test (for testing differences from random interactions) and from a Wilcoxon rank sum test (for comparison between conditions). **(H)** Same as (G), for enhancer-enhancer interactions between differentially acetylated TADs located within 20 Mb blocks.

We first focused on megabase-sized chromatin domains. We found that higher-order chromatin structures, including A/B compartments, TAD boundary location and strength, and CTCF loops, were largely unaffected by transient TPO signaling (Supplemental Fig. S4A-G). This result is in line with previous studies of extracellular signaling that examined higher-order chromatin architectures one hour after stimulation in other cellular systems (Jin et al. 2013; Le Dily et al. 2014).

We then analyzed the distribution of DARs within TADs identified in basal condition. This analysis revealed a clear spatial segregation of homotypic TPO-responsive DARs within topological domains (Fig. 3B), indicating that changes in cis-regulatory activity induced by TPO are spatially correlated at the level of TADs. To more formally investigate this aspect, we identified TADs exhibiting significantly altered H3K27ac levels in response to TPO at 1% FDR (Fig. 3C, see Methods). This analysis singled out 200 (6% of all H3K27ac-marked TADs) globally induced and 535 (16%) globally repressed TADs (henceforth collectively called differentially acetylated TADs), allowing us to study topological domains that were mostly perturbed by TPO.

Compared to TADs with no significant changes in H3K27 acetylation, differentially acetylated TADs were moderately enriched (approximately twofold) for differentially transcribed genes. As expected, genes in repressed TADs showed an overall lower expression than genes in activated TADs (Fig. 3D). Notably though, while only 3% of downregulated genes were located in activated TADs, a much higher fraction (12.9%) of upregulated genes lay within repressed TADs (Fig. 3E). Thus, whereas downregulated genes were virtually excluded from activated TADs, repressed TADs appeared at least partially permissive to transcriptional upregulation (Fig. 3E).

Finally, we examined the relation between transcriptional and cis-regulatory responses to TPO by correlating the number of differentially transcribed genes with the number of DARs located within the same differentially acetylated TAD. As expected, we found a marked correlation between increased cis-regulatory activity and transcriptional upregulation (Fig. 3F). In contrast, we noticed a weaker association between repressed cis-regulatory elements and transcriptionally downregulated genes (Fig. 3F). To further explore this aspect, we analyzed the distribution of differentially transcribed genes within all TADs exhibiting at least two homotypic DARs. To our surprise, we found that most TADs with repressed DARs did not show a corresponding detectable repression of genes located therein (Supplemental Fig. S4G), thus explaining the relatively weak association between cis-regulatory and transcriptional repression. Importantly, basal expression levels of these genes were significantly lower than for genes located within TADs hosting at least one differentially transcribed gene (Supplemental Fig. S4H). The lower expression levels reduce our ability to detect statistically significant further reductions in transcript levels. However, our results raise the possibility that repressed DARs within these loci might represent the first detectable step towards stable gene silencing of lineage-inappropriate genes.

### TPO-responsive enhancers are spatially clustered

At finer scales, physical interactions between promoter-distal sites appear to be widespread (Fullwood et al. 2009; Sahlén et al. 2015; Ghavi-Helm et al. 2014; Phillips-Cremins et al. 2013) and might function to provide specificity and robustness to enhancer-promoter interactions within cis-regulatory units (Ing-Simmons et al. 2015; Markenscoff-Papadimitriou et al. 2014). Previous work suggests that enhancer elements tend to cluster in the nuclear space in a cohesin-dependent manner (Ing-Simmons et al. 2015), but how enhancer-enhancer interactions are modulated by extracellular signaling remains largely unknown.

To investigate this aspect, we took advantage of the extensive and spatially compartmentalized epigenome remodeling induced by transient TPO signaling, and analyzed enhancer-enhancer interactions within and between differentially acetylated TADs using a structured interaction matrix analysis (SIMA) (Lin et al. 2012). This method pools Hi-C interactions across a pre-defined set of genomic regions (in our case, enhancer elements) and computes their interaction strength relative to control regions randomly sampled from the same set of chromatin domains (differentially acetylated TADs). First, we focused on enhancer-enhancer interactions within activated and repressed TADs. In the absence of TPO, we found that homotypic enhancers interacted significantly more frequently with each other than expected based on a null model (Fig. 3G), indicating that they tend to congregate within topological domains. Importantly, this was not the case when enhancer coordinates were systematically shifted by 10 kb along the chromatin fiber, demonstrating that our analysis is well calibrated (Fig. 3G). We then examined the consequences of TPO signaling. Interestingly, enhancer-enhancer interactions were only moderately perturbed within 30 minutes of TPO (Fig. 3G). Indeed, homotypic enhancers remained significantly clustered within differentially acetylated TADs, suggesting that TPO selectively modulates enhancer-enhancer interactions rather than altering them at a global scale.

Although by definition intra-TAD interactions occur more frequently than interactions spanning TAD boundaries, topological domains represent a modest twofold enrichment in interaction frequency (Dixon et al. 2012; Nora et al. 2012; Sexton et al. 2012). Therefore, we tested whether enhancers located within neighboring TPO-regulated TADs show evidence of spatial clustering (see Methods). We found that, similarly to intra-TAD interactions, inter-TAD enhancer-enhancer interactions were significantly enriched over random expectation and only moderately perturbed by transient TPO signaling (Fig. 3H). This enrichment was further confirmed by an analysis of enhancer-enhancer interactions between CTCF loops (Supplemental Fig. S3F).

Together, these results indicate that TPO-responsive enhancers engage in preferential long-range intra‐ and inter-TAD interactions resulting in their clustering in the three-dimensional nuclear space, and that TPO-induced changes in enhancer activity are at least partly uncoupled from this spatial clustering of enhancer elements.

### TPO-responsive super-enhancers control the expression of key hematopoietic regulators

Transcriptional enhancers not only interact with each other through long-range interactions but also within dense enhancer hotspots known as super-enhancers (SEs) or stretch-enhancers (Whyte et al. 2013; Parker et al. 2013; Ing-Simmons et al. 2015), which have been implicated in the transcriptional control of cell identity genes (Hnisz et al. 2013). Intriguingly, a recent study revealed that SE-associated genes are enriched for functional categories relevant to cytokine biology in mouse T lymphocytes (Vahedi et al. 2015). Therefore, we reasoned that SEs might be prime candidates for the integration of TPO signaling to chromatin.

To test this hypothesis, we identified SEs in serum-starved HPC-7 cells (Fig. 4A, see Methods) and analyzed their response to 30 minutes TPO stimulation. TPO signaling significantly perturbed the activity of more than half of all SEs (162 out of 277, 58%) identified in basal condition. Interestingly, the vast majority of these TPO-responsive SEs (140/162, 86%) were significantly deacetylated within 30 minutes of TPO, indicating that TPO signaling primarily represses SE activity (Fig. 4B and Supplemental Fig. S5A-B). We next examined the consequences of this perturbation on the transcription of putative target genes as a function of their genomic distance to SEs. Differential acetylation of SEs was more frequently linked to transcriptional perturbation of SE-proximal genes located within genomic distances <1 Mb (Fig. 4C and Supplemental Fig. S5C), with 27% of activated SEs and 22% of repressed SEs localizing within the same CTCF loop as their closest differentially transcribed gene. Nevertheless, the majority of genes located in close proximity to TPO-responsive SEs exhibited little or no detectable transcriptional changes within 30 minutes of TPO (Fig. 4C and Supplemental Fig. S5C). This result suggests that SE chromatin dynamics might often precede transcriptional changes at target genes or that genomic proximity is not an accurate predictor of SE targets.

**Figure 4.**
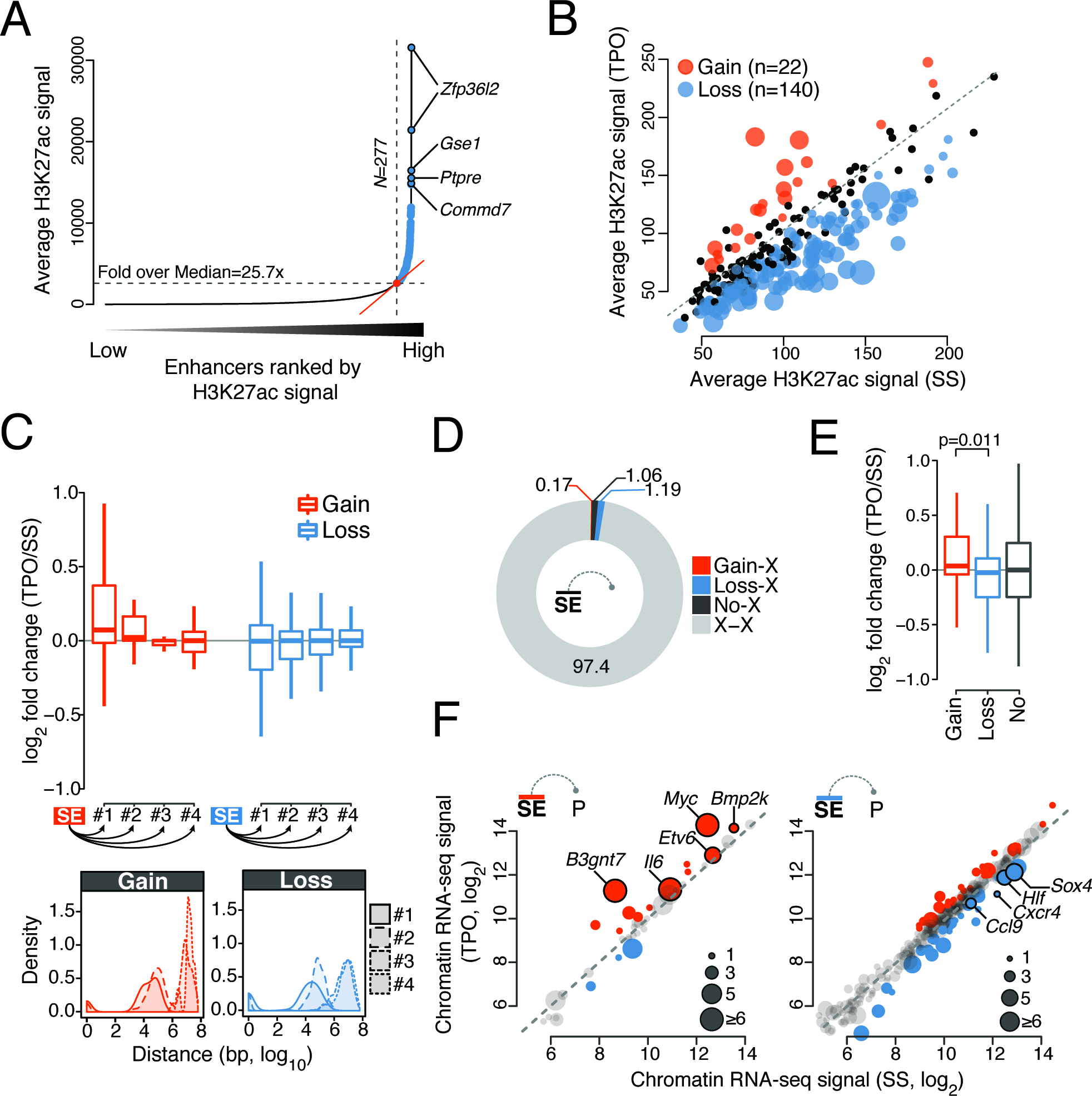
TPO signaling is integrated at super-enhancers. **(A)** Distribution of average (control-subtracted) normalized H3K27ac ChIP-seq signals at enhancer regions in serum starved HPC-7 cells. Super-enhancers (SEs, blue) exhibit exceptionally high H3K27ac levels. **(B)** Average normalized H3K27ac signals per kb of SE. Significantly differentially acetylated SEs are highlighted. Point sizes are proportional to ‐log_10_(q-value). **(C)** (top) Distribution of chromatin-associated RNA expression fold change for first, second, third and fourth nearest gene to a differentially acetylated SE. (bottom) Corresponding distribution of genomic distances. **(D)** Percentage of significant promoter Capture Hi-C interactions anchored at SEs. X, any HindIII restriction fragment located outside SEs. **(E)** Distribution of chromatin-associated RNA expression fold changes of SE-target genes defined by promoter Capture Hi-C. p-value is from a Wilcoxon rank sum test. **(F)** Chromatin-associated RNA expression (regularized log2 values) of gene targets of differentially acetylated SE as defined in (E). Point sizes are proportional to the number of constituent enhancers exhibiting significant interactions with the gene promoter. Selected SE targets are labeled.

To more accurately link SEs to their target genes, we enriched our Hi-C libraries for interactions anchored at 22,225 annotated gene promoters within the mouse genome using Promoter Capture Hi-C (PCHi-C) (Schoenfelder et al. 2015), for a total of 445 and 407 million intrachromosomal valid di-tags in basal and TPO-treated conditions, respectively. Data normalization and interaction detection from two biological replicates using the CHiCAGO pipeline (Cairns et al. 2016) resulted in 192,634 and 181,235 statistically significant (CHiCAGO score ≥ 5) promoter-anchored long-range cis-interactions for basal and TPO-treated conditions, respectively. Focusing on SEs, we found that approximately 2.4% of all significant PCHi-C interactions linked promoters to SE constituents (Fig. 4D), allowing us to infer the targets of differentially acetylated SEs based on spatial proximity. While most SE-interacting genes showed no detectable transcriptional changes within 30 minutes of TPO, we found that targets of induced and repressed SEs were enriched for transcriptionally up‐ and downregulated genes, respectively, and these included known SE targets such as *Myc* and *Etv6* (Khan and Zhang 2016) (Fig. 4E-F). Interestingly, promoters of key hematopoietic regulators rapidly repressed by TPO signaling (*Hlf*, *Sox4* and *Cxcr4*, Fig. 1B-C) were highly connected to constituent enhancers within repressed SEs (Fig. 4F), suggesting that their expression is controlled from a distance through TPO-responsive SEs.

### Rapid modulation of cis-regulatory activity is largely independent of chromatin looping dynamics

Next, we tested whether the pervasive epigenome remodeling observed within differentially acetylated TADs and within SEs was accompanied by systematic changes in chromatin looping. To this end, we computed normalized Hi-C contact frequencies within these chromatin domains before and after 30 minutes TPO stimulation. Our analysis revealed no systematic differences in contact frequencies between conditions, which exhibited nearly perfectly correlated Hi-C signals (r > 0.99) (Fig. 5A). This was further confirmed by a systematic analysis of chromatin interactions anchored at DARs (Fig. 5A). These results indicate that, despite substantial changes in both transcriptome and epigenome, chromatin architectures at differentially acetylated TADs and SEs were essentially unaltered following TPO stimulation.

**Figure 5.**
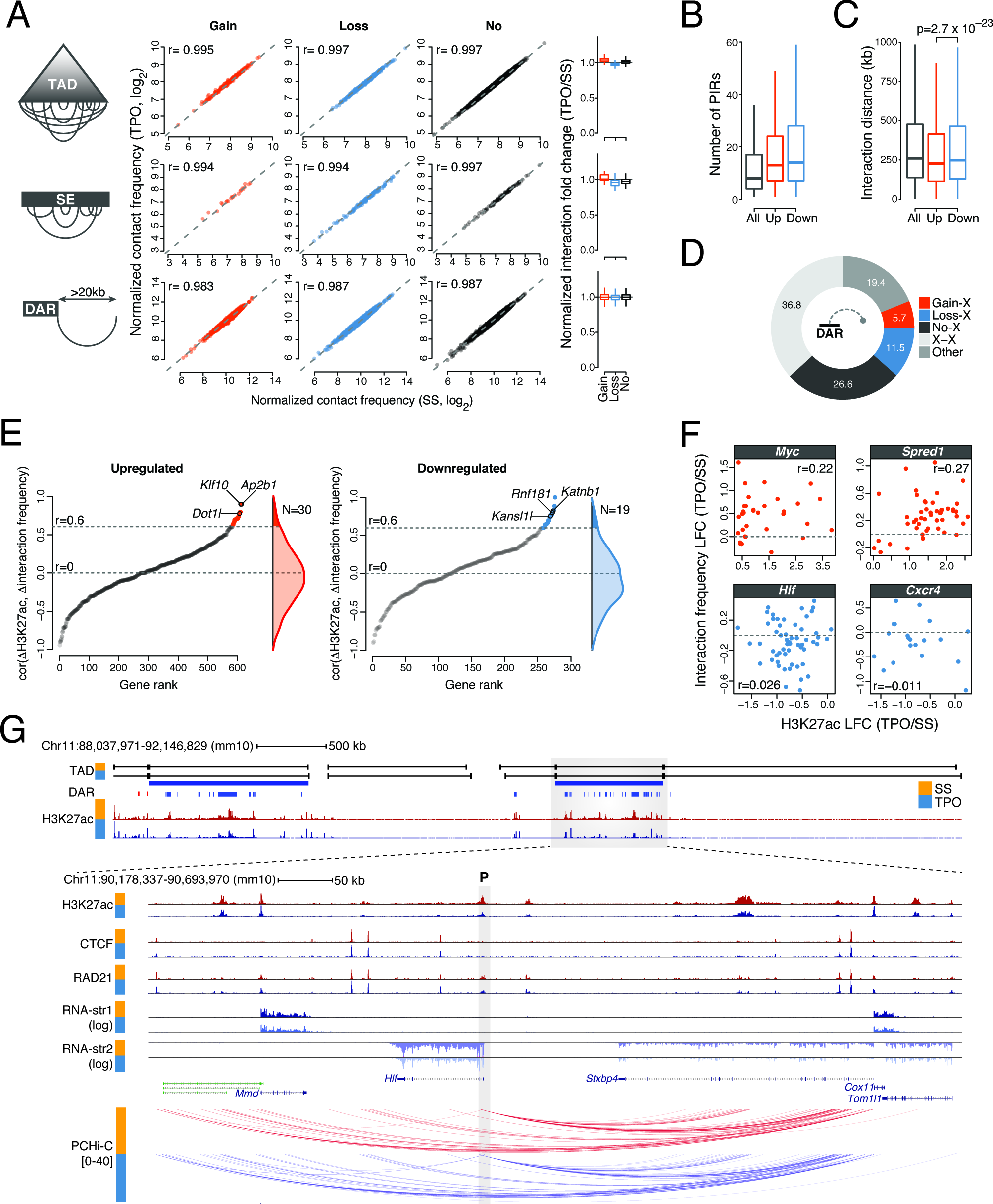
Rapid modulation of cis-regulatory activities within poised chromatin architectures. **(A)** Normalized Hi-C contact frequencies for intra-TAD and intra-SE interactions (per kb of element), and for interactions anchored at DARs spanning more than 20kb. Spearman’s rank correlation coefficients (r) are shown. Boxplots (right) summarize normalized interaction fold change distributions. **(B)** Distribution of the number of promoter-interacting regions (PIRs) for all baited promoters (All) and promoters of transcriptionally up‐ and down-regulated genes. **(C)** Distribution of interaction distances for all baited promoters (All) and promoters of transcriptionally up‐ and down-regulated genes. p-value is from a Wilcoxon rank sum test. **(D)** Percentage of significant promoter Capture Hi-C interactions anchored at DARs. X, any HindIII restriction fragment located outside DARs. **(E)** Relation between cis-regulatory activity and chromatin architectures at cis-regulatory units for transcriptionally up‐ and down-regulated genes (see Methods). Genes are ranked based on the Spearman’s rank correlation coefficient (r) between normalized H3K27ac fold change (FC, TPO/SS) and normalized Capture Hi-C interaction fold change at target DARs. Only promoters exhibiting significant interactions with at least five distinct DAR-containing HindIII restriction fragments were considered. Genes exhibiting significant correlations are colored, and representative hits are labeled. **(F)** Representative examples from the analysis in (E). Each dot corresponds to an HindIII restriction fragment. LFC, log2 fold change. **(G)** Epigenomic configuration of the *Hlf* locus. Statistically significant promoter Capture Hi-C interactions within the *Hlf* TAD are shown. The gray shaded rectangle denotes the position of the baited *Hlf* promoter (P). str, strand.

We then focused on the spatial chromatin architecture at differentially transcribed gene loci, taking advantage of the increased resolution offered by PCHi-C over Hi-C data. We found that most promoters of differentially transcribed genes engaged > 10 PIRs (with a median of 13 and 14 PIRs for transcriptionally up‐ and down-regulated genes, respectively) (Fig. 5B) and although for transcriptionally upregulated genes these interactions tended to span significantly shorter genomic distances (Fig. 5C), no major differences were found in the connectivity of these loci. In addition, 63% of all promoter interactions detected by PCHi-C were anchored at H3K27ac-marked regions (Fig. 5D), with 30.7% encompassing at least one DAR, indicating extensive coverage of these genomic regions.

These results prompted us to systematically correlate changes in H3K27ac levels with changes in PCHi-C interaction frequency within individual differentially transcribed cis-regulatory units (see Methods). For each promoter, we defined a cis-regulatory unit as the union of all its H3K27ac-marked PIRs, and we set a threshold of at least five such regions being required to ensure robust estimates of H3K27ac fold changes and correlation coefficients. This resulted in a set of 907 differentially transcribed loci (Fig. 5E). Interestingly, we found that the vast majority of cis-regulatory units exhibited little or no correlation between changes in cis-regulatory activity and chromatin looping (Fig. 5E).

This result was further supported by a permutation test where the connectivity of cis-regulatory units was randomly scrambled to derive a null distribution for the correlation coefficients (see Methods). Our data revealed that only 5.4% of differentially transcribed gene loci being tested (49 out of 907, corresponding to 4.8% and 6.7% of analyzed transcriptionally up‐ and down-regulated genes) exhibited a statistically significant correlation between acetylation and looping dynamics in response to transient TPO signaling (Fig. 5E). Neither rapidly induced genes (e.g. *Myc* and *Spred1*, Fig 1A) nor repressed loci encoding key hematopoietic regulators belonged to this set (Fig. 5F). Indeed, while these loci exhibited extensive differential acetylation within 30 minutes of TPO, this was not accompanied by a rewiring of chromatin loops (Fig. 5F-G and Supplemental Fig. S6).

Together, these results demonstrate that the extensive and spatially compartmentalized epigenome remodeling induced by transient TPO signaling takes place within poised chromatin architectures, and suggest that the rapid cytokine-dependent modulation of cis-regulatory activity is largely independent of chromatin looping dynamics.

### TF binding patterns but not DNA sequence features accurately predict rapid cis-regulatory responses to TPO

Finally, we asked which TFs orchestrate the rapid cis-regulatory dynamics induced by TPO signaling. To identify TFs that control activated and repressed cis-regulatory elements within DARs, we subjected differentially acetylated DHSs to a de novo motif discovery analysis. Interestingly, we found that both activated and repressed DARs were strongly enriched for ETS motifs, which feature a central 5’-GGAA-3’ core, and for motifs recognized by the RUNX and GATA families of TFs (Fig. 6A-B). These motifs exhibited similar enrichments and overlapping distributions at activated and repressed DARs (Fig. 6B). In stark contrast, an interferon g-activation sequence (GAS) motif, which is recognized by tyrosine-phosphorylated STAT (pSTAT) TFs, was specifically overrepresented at activated but not at repressed DARs (Fig. 6B), indicating that canonical JAK/STAT signaling is exclusively associated with DAR activation. Surprisingly, however, a systematic motif enrichment analysis based on 363 position weight matrices (PWMs) for 322 vertebrate TFs revealed that no TF motif other than those recognized by pSTATs exhibited a comparably specific association with either DAR class (Fig. 6C). Thus, despite exhibiting opposite response to TPO, activated and repressed cis-regulatory elements were characterized by a remarkably similar TF motifs composition.

**Figure 6.**
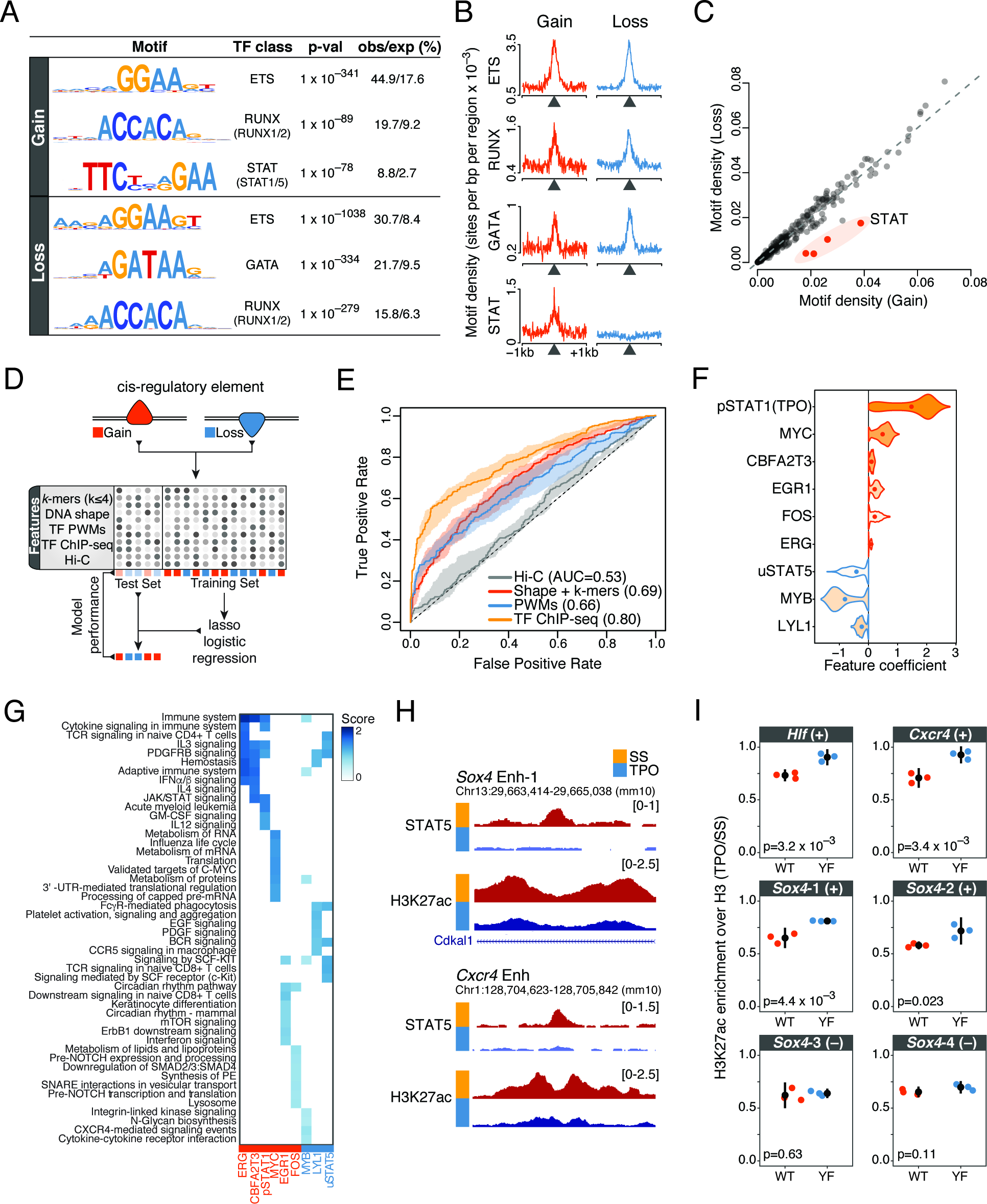
TF binding patterns accurately predict rapid cis-regulatory responses to TPO. **(A)** Top transcription factor motifs (position weight matrices, PWMs) identified by de novo motif discovery analysis within 200 nt of the summit of differentially acetylated DHSs (see Methods). The percentage of observed and expected motif occurrence is indicated. **(B)** Motif density (PWMs) for the indicated transcription factor motifs within ±1 kb of the summit of differentially acetylated DHSs. **(C)** Total motif density within 200 nt of the summit of differentially acetylated DHSs, for a collection of 363 vertebrate transcription factor PWMs. **(D)** Schematic outline of the statistical learning strategy used to predict how cis-regulatory elements within DARs respond to TPO. **(E)** Test set receiver operating characteristic (ROC) curve and area under the ROC curve (AUC) values for lasso models trained on the indicated sets of features. The shaded area is delimited by the ROC curves for models with highest and lowest AUC values, whereas the ROC curve for the model closest to mean AUC is shown. **(F)** Top-ranked features selected by bootstrap-lasso for the model trained on TF ChIP-seq profiles, ranked by selection probability (stability). Violin plot lines are color-coded according to coefficient signs (positive, red; negative, blue). **(G)** Genomic regions enrichment of annotations tool (GREAT) analysis of differentially acetylated cis-regulatory elements bound by TFs in (F). The top eight significantly enriched Molecular Signature Database (MSigDB) pathways were considered for each transcription factor, ranked by binomial p-value. **(H)** Representative tracks illustrating STAT5 binding dynamics at differentially acetylated enhancers within key hematopoietic gene loci. **(I)** Relative H3K27ac enrichment (TPO/SS, normalized to total H3 levels) at the indicated enhancer elements (uSTAT5-bound (+) or uSTAT5 negative (-, control)) for HPC-7 cells expressing wild-type (WT) or mutant (Y699F) STAT5B. Tested loci include the enhancers shown in (H). Error bars are mean±SD (n=3). p-values are from a two-sided Welch’s *t*-test.

This result led us to perform an unbiased survey of DNA and chromatin features that might be able to predict rapid cis-regulatory responses to TPO signaling. To this end, we considered activated and repressed cis-regulatory elements and trained least absolute shrinkage and selection operator (lasso) logistic regression models (Tibshirani 1996) using different sets of features (Fig. 6D) (see Methods). These included DNA sequence *k*-mers (2 ≤ *k* ≤ 4), DNA shape features (Chiu et al. 2016), a collection of > 1,700 single and composite TF motifs (Diaferia et al. 2016) that was filtered for TFs expressed in HPC-7 cells, a collection of 29 ChIP-seq binding profiles for hematopoietic and other sequence-specific TFs in HPC-7 cells before (n = 26) and after (n = 3) 30 minutes TPO stimulation (generated as part of this study or previously published (Wilson et al. 2010, 2016; Park et al. 2015), see Supplemental Table S1), and normalized Hi-C signals as a proxy for interaction frequencies. Model performances were evaluated on a test set consisting of differentially acetylated cis-regulatory elements that were not used for learning model parameters (Fig. 6D).

Interestingly, while DNA sequence features were only moderately predictive for cis-regulatory responses to TPO (area under the receiver operating characteristic curve, AUC < 0.70), in vivo TF binding profiles were able to accurately predict changes in cis-regulatory activity (AUC = 0.80) (Fig. 6E). In contrast, Hi-C signals showed virtually no predictive power (AUC = 0.53) (Fig. 6E), further supporting the notion that these changes occur within poised chromatin architectures. In addition, more complex models combining sequence and chromatin features did not outperform models learnt on TF binding profiles alone (Supplemental Fig. S7A), suggesting that key TFs were already included within our chromatin feature set. Furthermore, these results were validated by an independent learning scheme using random forest classifiers (Breiman 2001) (Supplemental Fig. S7B).

Next, we focused on the models learnt on TF binding profiles and sought to identify the chromatin features that contribute the most to model predictions by estimating feature stability coefficients using bootstrap-lasso (Comoglio and Paro 2014; Comoglio et al. 2015). Intuitively, the more a feature is necessary for accurate predictions, the higher its stability value. Our analysis revealed that pre-existing MYC binding and binding of pSTAT1 upon TPO stimulation were stably associated with the activation of cis-regulatory elements (Fig. 6F and Supplemental Fig. S7C). In contrast, binding of MYB, LYL1 and tyrosine unphosphorylated STAT5 (uSTAT5) in the absence of TPO was predictive for repression of cis-regulatory elements (Fig. 6F and Supplemental Fig. S7C). Notably, DARs bound by stably selected features were significantly associated with distinct signaling pathways and metabolic functions, spanning the whole spectrum of biological processes regulated by TPO (Fig. 6G).

We previously demonstrated that in the absence of TPO, chromatin-bound uSTAT5 restrains a megakaryocytic transcriptional program in hematopoietic stem/progenitor cells, and that TPO-mediated phosphorylation of STAT5 triggers a genome-wide relocation of STAT5 to canonical GAS motifs (Park et al. 2015). Intriguingly, uSTAT5 knockdown not only upregulated megakaryocytic-affiliated genes, but it also repressed lineage-inappropriate genes including lymphoid‐ and pre-granulocyte/monocyte-affiliated genes (Park et al. 2015). This result, along with our feature importance analysis (Fig. 6F), led us to hypothesize that uSTAT5 binding might be required to maintain the activity of cis-regulatory elements within gene loci rapidly repressed by TPO. To test this hypothesis, we took advantage of a mutant (Y699F) STAT5B that ablates TPO-induced tyrosine phosphorylation of STAT5 and does not act as dominant negative (Park et al. 2015). We expressed wild-type or mutant STAT5B in HPC-7 cells, and measured H3K27ac levels at uSTAT5-bound enhancers within the *Hlf*, *Sox4* and *Cxcr4* loci (Fig. 6H) by ChIP-qPCR before and after 30 minutes TPO stimulation. Interestingly, the Y699F mutant significantly attenuated deacetylation at uSTAT5-bound enhancers in response to TPO compared to wild-type (Fig. 6I), indicating that uSTAT5 binding is required to maintain the activity of these elements.

## Discussion

In this study, we focus on the immediate early consequences of TPO signaling to chromatin and examine the relation between signal-dependent modulation of cis-regulatory activity and chromatin looping. Our work provides three key findings.

First, we present evidence for rapid and pervasive changes in cis-regulatory activity that are readily detectable within 30 minutes of TPO. Interestingly, this widespread epigenome remodeling encompasses up to 40% of active genomic regions marked by H3K27ac prior to TPO stimulation, indicating that short-term cytokine signaling is sufficient to profoundly alter chromatin states at cis-regulatory elements. Notably, a sizeable fraction of these remodeling events corresponds to histone deacetylation of enhancer elements associated with genes playing key roles in innate and adaptive immune cells. This rapid deacetylation of enhancer elements likely represents the first step towards decommissioning of active enhancers associated with lineage-inappropriate transcriptional programs (Smith and Shilatifard 2014), and suggests that TPO signaling rapidly dismantles cis-regulatory networks affiliated to alternative lineages.

Secondly, we demonstrate that TPO-induced modulation of cis-regulatory units occur largely independently of alterations in chromatin looping, indicating that a responsiveness to TPO is hardwired into a poised HSPC spatial genome architecture prior to cytokine exposure. It is important to note, however, that TPO-dependent modulation of short-range interactions between enhancers and promoters would not be detectable in our PCHi-C data.

A previous Hi-C analysis of chromatin structures after 1h TNF stimulation of primary human fibroblast cells showed that TNF-responsive enhancers are already in contact with their target promoters prior to stimulation, and these pre-existing chromatin structures appear to be strong predictor of gene induction (Jin et al. 2013). Similarly, pre-formed chromatin loops have also been detected at inducible gene loci regulated by TP53, FOXO3 and hormone receptor binding by 3C or circular chromosome conformation capture (4C) sequencing (Melo et al. 2013; Eijkelenboom et al. 2014; Jin et al. 2013). Our results indicate that poised chromatin architectures not only provide permissive regulatory topologies for transcriptional induction, but also a platform for rapid signaling-dependent repression of cis-regulatory units. Therefore, it is plausible that disruption of pre-formed chromatin architectures prior to TPO stimulation might similarly impair both activation and repression of cis-regulatory units in response to TPO signaling. However, further work will be required to formally test this hypothesis.

Thirdly, we show that TPO signaling can induce opposite responses at cis-regulatory elements exhibiting markedly similar DNA sequence compositions and regulatory codes. This unexpected finding raises the question about which molecular determinants direct these opposing events. Regulatory instructions are encoded in DNA sequences by the identity, frequency, affinity and grammar of TF binding sites. However, how these parameters dictate cis-regulatory dynamics is poorly understood (Levo and Segal 2014; Spitz and Furlong 2012). Our results indicate that, with the notable exception of pSTATs, the mere presence or absence of binding motifs for specific TFs cannot predict individual cis-regulatory responses to TPO and raise the possibility that orientation and spacing of TF motifs might play a role in determining these responses. Consistent with this concept, a recent work demonstrated that a different motif grammar based on the same building blocks defines activating and repressing cis-regulatory elements in rod photoreceptors (White et al. 2016). Moreover, several studies showed that a subset of TFs can act both as repressor or activator depending on cellular states, sequence contexts and binding of co-factors (Methot and Basler 1999; Sharrocks 2001; Hollenhorst et al. 2011; Nayak et al. 2009; Liu et al. 2014; Sanchez-Tillo et al. 2015). Deciphering the motif grammar that distinguishes activated and repressed cis-regulatory elements should be an important goal of future studies. To this end, statistical learning models taking into account motif orientation and spacing could be used to guide the design of massively parallel reporter assays based on synthetic regulatory elements or more targeted in vitro assays (Fiore and Cohen 2016; Inoue and Ahituv 2015).

While our analysis suggests that regulatory priming in multipotent progenitors does not appear to be constrained by sequence composition, in vivo TF binding patterns are able to accurately predict rapid cis-regulatory responses to TPO. Interestingly, feature importance analysis indicates that pre-existing TF binding prior to TPO stimulation significantly contributes to these predictions. This notion is further supported by the targeted experimental analysis of uSTAT5-bound enhancers where pre-existing uSTAT5 binding is predictive for the repression of these regulatory elements. Moreover, our statistical models identified other potential key players mediating rapid cis-regulatory responses to TPO such as MYB and LYL1. These hematopoietic TFs are critically involved in the maintenance of HSPCs as well as in lymphoid differentiation (Lieu and Reddy 2009; Fahl et al. 2009; Greig et al. 2010; Souroullas et al. 2009; Capron et al. 2006). Our results suggest that MYB and LYL1 binding prior to TPO stimulation predicts deacetylation of cis-regulatory elements associated with the lymphoid lineage. Therefore, it is tempting to speculate that TPO supports megakaryocytic lineage specification in part by restricting the developmental potential of multipotent progenitor cells. To this end, it would be interesting to further investigate whether TPO directly regulates chromatin binding and/or activity of MYB and LYL1 in HSPCs.

Taken together, our study unravels the multifaceted immediate early consequences of TPO signaling to chromatin and provides a paradigm for the integrative epigenomic analysis of cytokine signaling.

## Methods

### Cells and cell culture

HPC-7 cells were grown at 37°C and 5% CO2 in IMDM (Invitrogen) supplemented with 10% fetal calf serum, 10% SCF conditioned media (produced by the BHK/MKL cell line), 1% L-Glutamine, 1% penicillin/streptomycin and 74.8μM monothioglycerol (Sigma). Cell density was maintained between 5×10^5^ - 2×10^6^ cells/mL.

### Mice

Eight to twelve weeks old C57BL/6 mice (Charles River) were used in this study. All mouse procedures were approved by the UK Home Office and the University of Cambridge Animal Welfare and Ethical Review Board.

#### Serum starvation and cytokine stimulation

Cells were spun down at 1,000 rpm for 5 min and washed once in PBS. The cell pellet was resuspended at a density of 1×10^6^ cells/mL in StemSpan SFEM medium (StemCell Technologies) and incubated for four hours. Serum-starved cells were then stimulated with recombinant murine thrombopoietin (TPO, Peprotech) at 100 ng/mL for 30 min. TPO was diluted in a small volume of fresh StemSpan SFEM medium prior to stimulation.

#### Isolation of CD41^+^LSK cells

Freshly isolated mouse bone marrow cells were stained with AF700-conjugated lineage cocktail (133313), APC-Cy7-conjugated anti-cKit (105826), Bv605-conjugated anti-Sca1 (108133) and FITC-conjugated anti-CD41 (133903). 7-AAD (Invitrogen) was used to exclude dead cells. All antibodies were purchased from BioLegend unless otherwise indicated. Cells were sorted on fluorescence-activated cell sorting (FACS) Aria (BD Bioscience) directly into StemSpan SFEM medium (StemCell Technologies). The sorted cells were incubated for 1 hour at 37°C and 5% CO2 prior to stimulation with TPO as described above.

#### HDAC inhibitor treatment

HPC-7 cells were starved as described above. Cells were pre-treated with HDAC inhibitors, 1μM of Trichostatin A or 5nM of Romidepsin (Selleckchem) during the final 30 min of serum starvation and stimulated with TPO at 100 ng/mL for 30 min.

#### STAT5 expression

Retrovirus was produced using Phoenix cells (ATCC). HPC-7 cells were infected with retrovirus expressing STAT5B or mutant (Y699F) STAT5B in the presence 8μg/mL polybrene (Millipore). After 24 hours, GFP positive cells were FACS sorted and cultured in IMDM media (Invitrogen) supplemented with 10% fetal bovine serum and 10% SCF conditioned media.

#### RNA preparation and RT-qPCR

Total RNA was isolated using Direct-zol (Zymo Research) and reverse transcribed into cDNA using SuperScript IV First Strand Synthesis System (Invitrogen) following the manufacturer instructions. Quantitative RT-PCR was carried out using SYBR Green Brilliant II low Rox and a Stratagene Mx3000P machine (Agilent Technologies). For primer sequences, see Supplemental Table S2.

#### Subcellular RNA isolation

Subcellular fractionation and RNA preparation were performed essentially as described (Bhatt et al. 2012) with modifications detailed in Supplementary Methods.

#### Chromatin immunoprecipitation

Chromatin immunoprecipitation (ChIP) assays on serum-starved and TPO stimulated HPC-7 cells were performed as previously described (Wilson et al. 2010) (see Supplementary Methods).

#### Hi-C

Hi-C libraries were generated essentially as described (Schoenfelder et al. 2015) with modifications detailed in Supplementary Methods.

#### Promoter Capture Hi-C

Promoter Capture Hi-C libraries were generated essentially as described (Schoenfelder et al. 2015) with modifications detailed in Supplementary Methods.

#### Computational analyses

Subcellular RNA-seq, ChIP-seq, Hi-C and promoter Capture Hi-C data processing and all computational analyses are detailed in Supplementary Methods.

## Data Access

RNA-seq, ChIP-seq, Hi-C and promoter Capture Hi-C raw and processed data from this study have been submitted to the NCBI Gene Expression Omnibus (GEO; http://www.ncbi.nlm.nih.gov/geo/) under accession number GSE100835.

## Acknowledgments

We would like to thank Bertie Göttgens for providing access to raw data prior to their publication, and David Flores-Santa-Cruz and Rebecca Hannah for technical support. We are grateful to Gioacchino Natoli, Chiara Balestrieri and Liv Austenaa (European Institute of Oncology) for help with subcellular RNA isolation and TF motif analysis, and to Luca Giorgetti and Zhan Yinxiu (Friedrich Miescher Institute) for discussion. F.C. was supported by an EMBO long-term fellowship (1305-2015 and Marie Curie Actions LTFCOFUND2013/GA-2013-609409). Work in the Fraser lab was supported by the UK Biotechnology and Biological Sciences Research Council (grant ref. BB/J004480/1). Work in the Green laboratory was supported by Bloodwise (grant ref. 13003), the Wellcome Trust (grant ref. 104710/Z/14/Z), the Medical Research Council, the Kay Kendall Leukaemia Fund, the Cambridge NIHR Biomedical Research Center, the Cambridge Experimental Cancer Medicine Centre, the Leukemia and Lymphoma Society of America (grant ref. 07037), and a core support grant from the Wellcome Trust and MRC.

## Author contributions

Conceptualization, F.C. and H.J.P; Methodology, F.C., H.J.P and S.S.; Formal Analysis, F.C., I.B. and D.B.; Investigation, F.C., H.J.P and S.S.; Resources, P.F. and A.R.G.; Writing-Original Draft, F.C.; Writing - Review & Editing, F.C., H.J.P, S.S., I.B., P.F. and A.R.G.; Visualization: F.C.; Supervision: S.S., P.F. and A.R.G.; Funding Acquisition, P.F. and A.R.G.

---

**Supplemental Figure S1.**
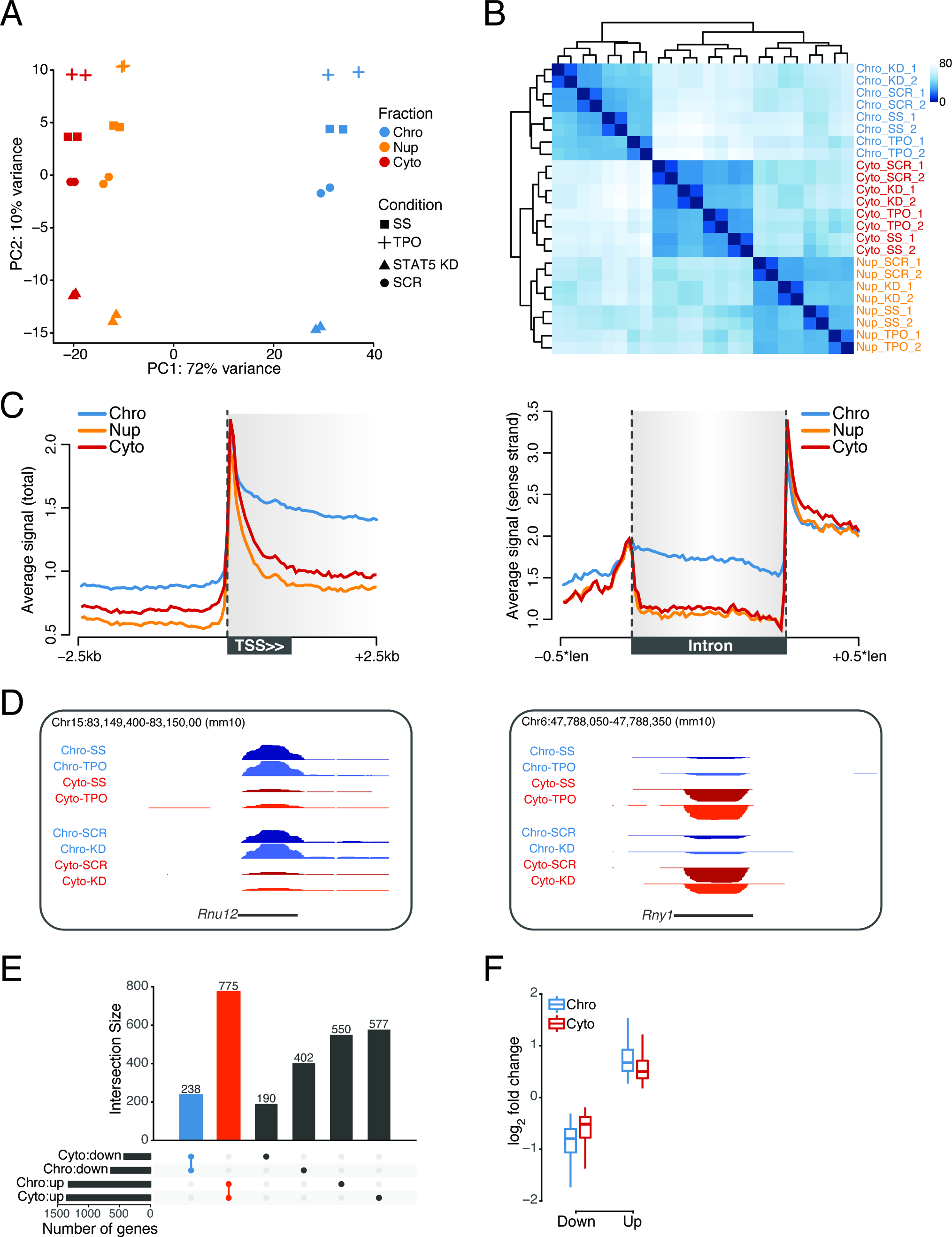
**(A)** Principal component analysis of gene expression profiles (regularized log_2_ expression values) measured by subcellular RNA-seq. Only genes with non-zero counts across all samples were considered. The percentage of variance explained by the first two principal components (PC) is indicated. SS, serum-starved (basal condition); KD, knockdown; SCR, scrambled shRNA control. **(B)** Hierarchical clustering of subcellular RNA-seq samples based on Euclidean distance of gene expression profiles. Only genes with non-zero counts across all samples were considered. **(C)** Average normalized chromatin-associated (Chro), nucleoplasmic (Nup) and cytoplasmic (Cyto) RNA-seq signal at isolated TSSs and introns exhibiting no feature overlap within the indicated genomic windows. **(D)** Representative tracks of chromatin-associated (Chro) and cytoplasmic (Cyto) RNA-seq signals at small nuclear and small cytoplasmic RNA loci (*Rnu12*, U12 small nuclear RNA; *Rny1*, Y1 small cytoplasmic RNA). **(E)** Summary of transcriptional changes induced by 30 minutes TPO stimulation of HPC7 cells. The intersection between differentially transcribed (chromatin-associated RNA-seq) and differentially expressed (cytoplasmic RNA-seq) genes is shown. **(F)** log_2_ fold change distribution of differentially transcribed (chromatin-associated RNA-seq) and differentially expressed (cytoplasmic RNA-seq) genes.

**Supplemental Figure S2.**
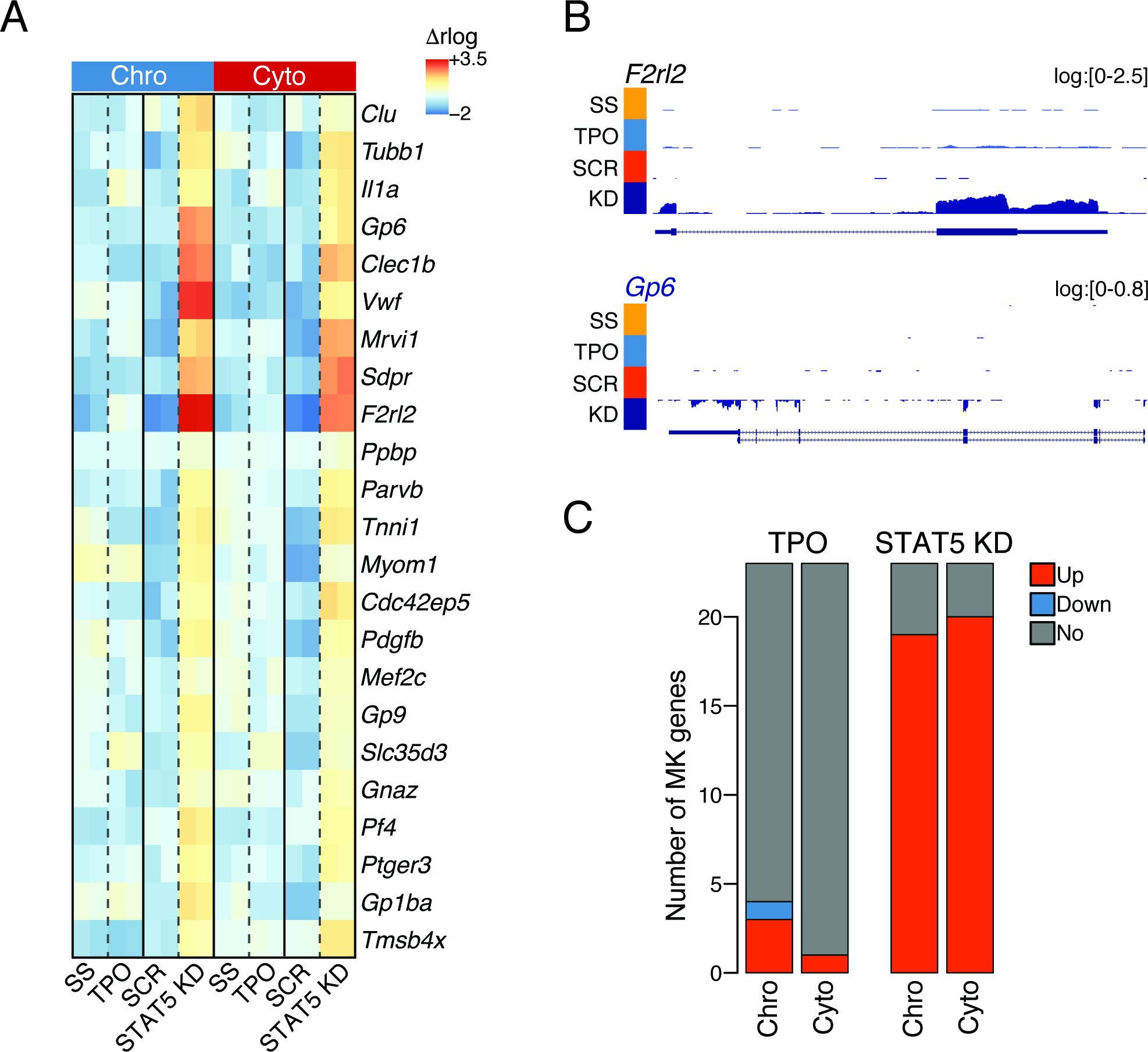
**(A)** Chromatin-associated (Chro) and cytoplasmic (Cyto) RNA expression heatmap for megakaryocytic genes upregulated upon shRNA-mediated STAT5 knockdown (KD) in HPC7 cells (Park et al. 2015), for the indicated conditions. Regularized log_2_ (rlog) expression values are row-mean subtracted. SCR, scrambled shRNA control. **(B)** Representative tracks of chromatin-associated RNA expression at megakaryocytic genes. RNA-seq coverage was log transformed with a pseudocount of 1. **(C)** Summary of megakaryocytic gene expression changes induced by 30 minutes TPO stimulation, compared to STAT5 knockdown (KD), from chromatin-associated (Chro) and cytoplasmic (Cyto) RNA fractions.

**Supplemental Figure S3.**
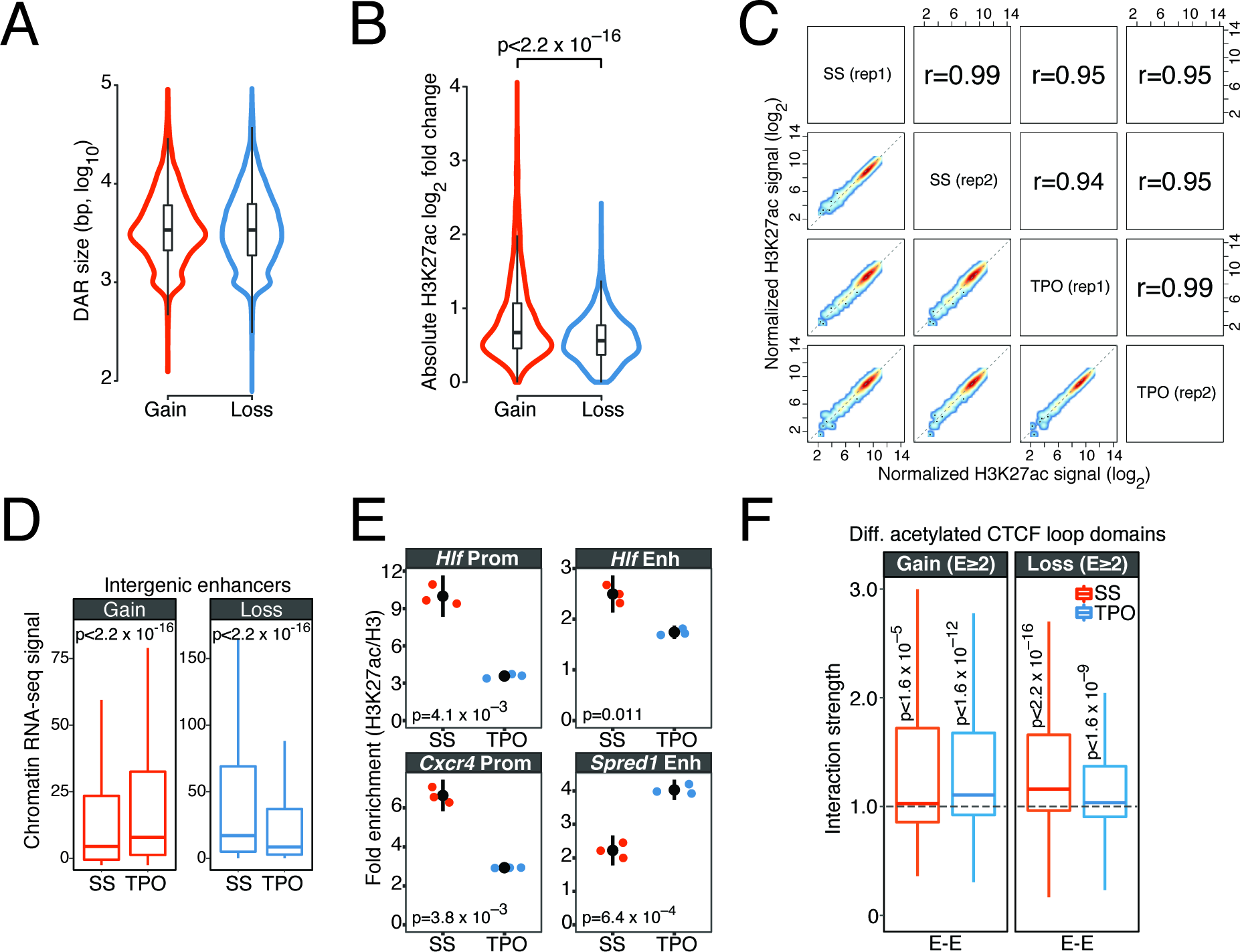
**(A)** Size distribution of activated (Gain) and repressed (Loss) DARs. **(B)** H3K27ac log_2_-fold change distribution for activated (Gain) and repressed (Loss) DARs. **(C)** Normalized H3K27ac signals within 5 kb windows centered on the transcription start site of the 1,000 most expressed genes in basal condition. Only genes that are not significantly differentially transcribed upon TPO were considered. Spearman’s rank correlation coefficients (r) are shown. **(D)** Normalized chromatin-associated RNA expression at differentially acetylated intergenic enhancers. p-values are from a Wilcoxon rank sum test. **(E)** H3K27ac levels normalized to total H3 occupancy at the indicated cis-regulatory elements (see Supplemental Table S2), measured by ChIP-qPCR. Error bars are mean±SD (n=3). p-values are from a two-sided Welch’s t-test. **(F)** Structured interaction matrix analysis (SIMA) of enhancer-enhancer interactions between CTCF loops located within 20 Mb blocks. The interaction strength reflects the enrichment of Hi-C interactions relative to randomly sampled genomic regions. p-values are from a Wilcoxon signed rank test.

**Supplemental Figure S4.**
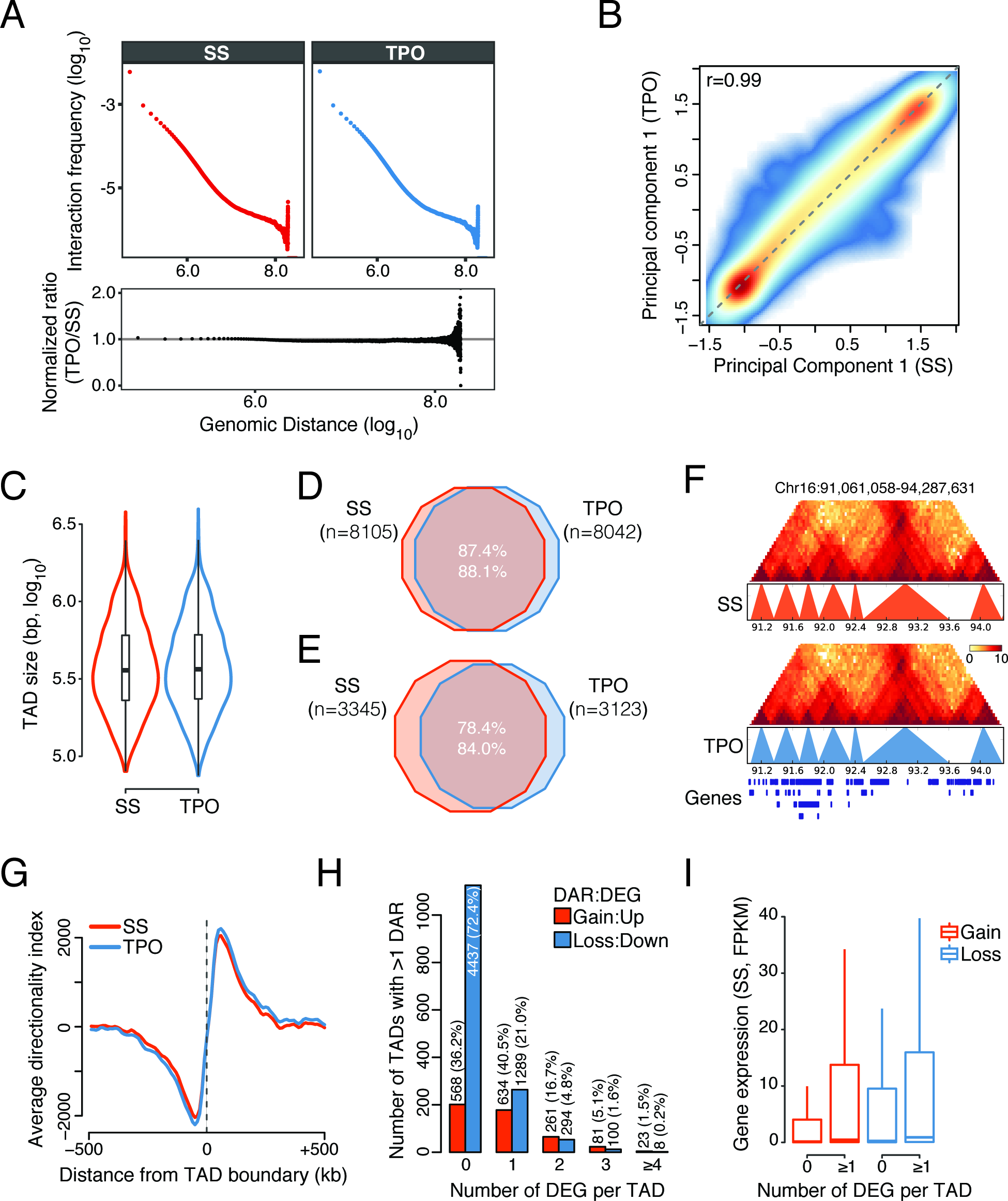
**(A)** Intrachromosomal interaction frequency as a function of genomic distance (top) and corresponding normalized interaction frequency ratio between conditions (bottom). **(B)** Genome-wide comparison of PC1 values defining A/B compartments identified in basal condition (SS) and after 30 minutes TPO stimulation. Principal component analysis was performed on normalized Hi-C contact matrices binned at 40 kb resolution. The Spearman’s rank correlation coefficient (r) is shown. **(C)** Size distribution of all detected TADs. **(D)** TAD boundary overlap (maximum tolerance of 10 kb) for all detected TADs. **(E)** Overlap between CTCF loops detected in basal condition (SS) and after 30 minutes TPO stimulation (maximum tolerance of 10 kb). **(F)** A representative 3.2 Mb genomic region (containing *Runx1*) depicting ICE (Iterative Correction and Eigenvector decomposition) normalized Hi-C signals at 40 kb resolution and TAD calls in basal condition (SS) and after 30 minutes TPO stimulation. **(G)** Average directionality index within ±500 kb of a TAD boundary. **(H)** Number of TADs with at least two DARs, partitioned by the number of differentially transcribed genes (DEG) therein. The total number and percentage of DARs within this set of TADs is indicated. **(I)** Distribution of basal gene expression levels (cytoplasmic RNA-seq) of genes located within the TADs in (H). FPKM, fragments per kb of exon per million reads mapped.

**Supplemental Figure S5.**
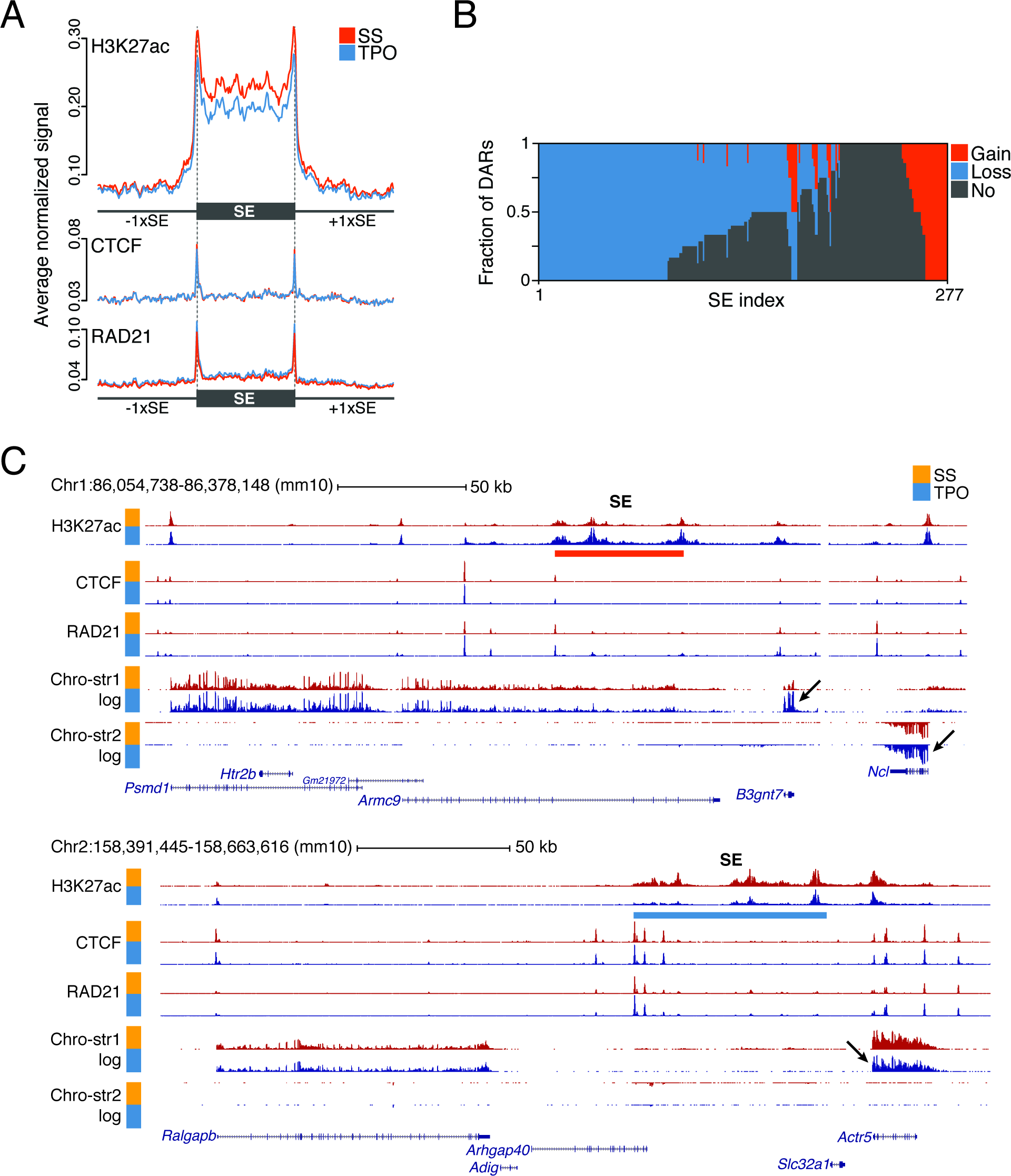
**(A)** Average normalized H3K27ac, CTCF and RAD21 ChIP-seq signals at SEs, including flanking regions of size equal to the SE length. Each genomic window was divided into 300 bins. **(B)** Distribution of DARs within SE. Each SE corresponds to a vertical line. (C) Representative tracks of differentially transcribed SE-proximal genes. RNA-seq coverage was log transformed with a pseudocount of 1.

**Supplemental Figure S6.**
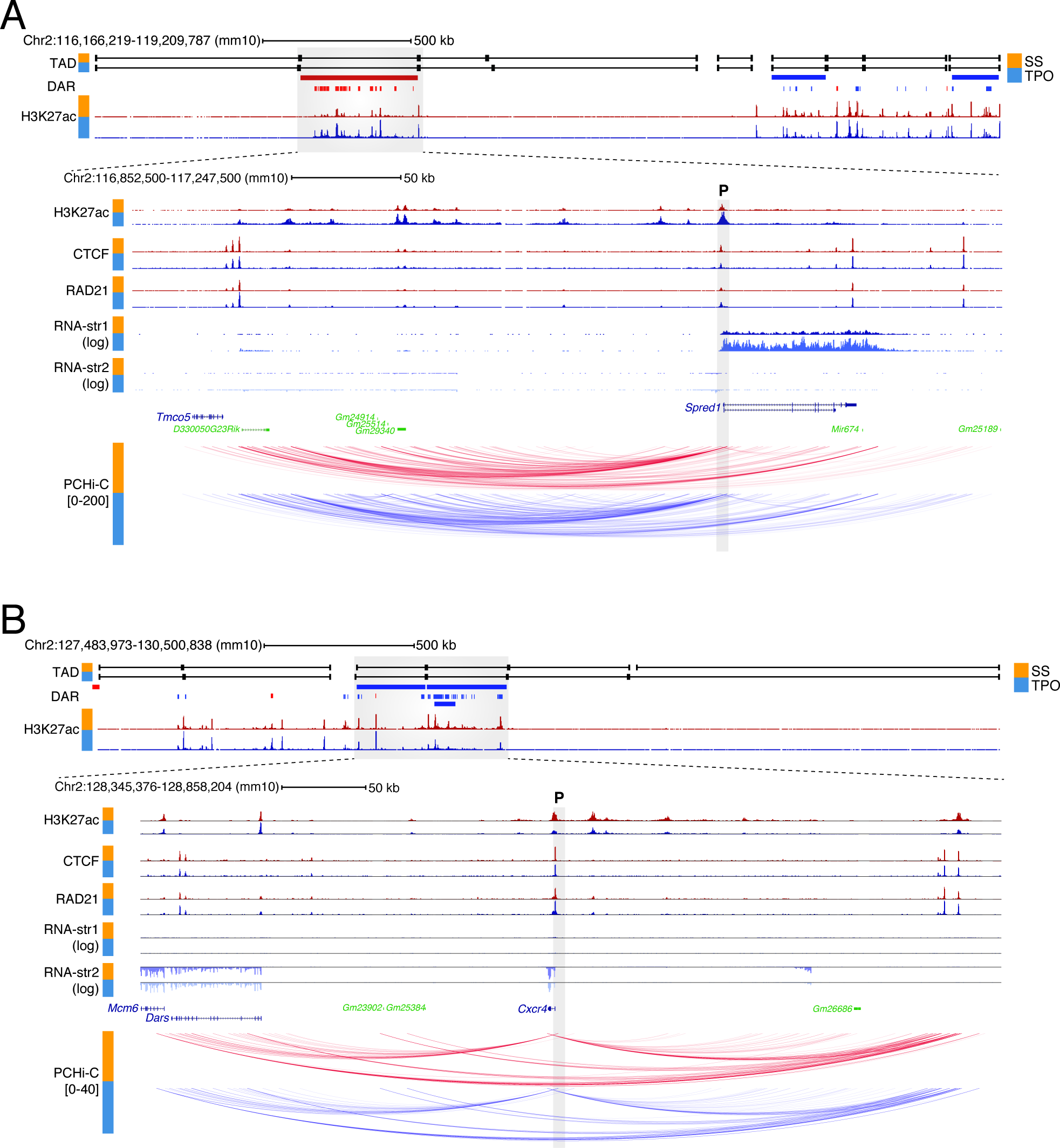
**(A)** Epigenomic configuration of the *Spred1* locus. Statistically significant promoter Capture Hi-C interactions within the *Spred1* TAD are shown. The gray shaded rectangle denotes the position of the baited *Spred1* promoter (P). str, strand. **(B)** Same as (A), for the *Cxcr4* locus.

**Supplemental Figure S7.**
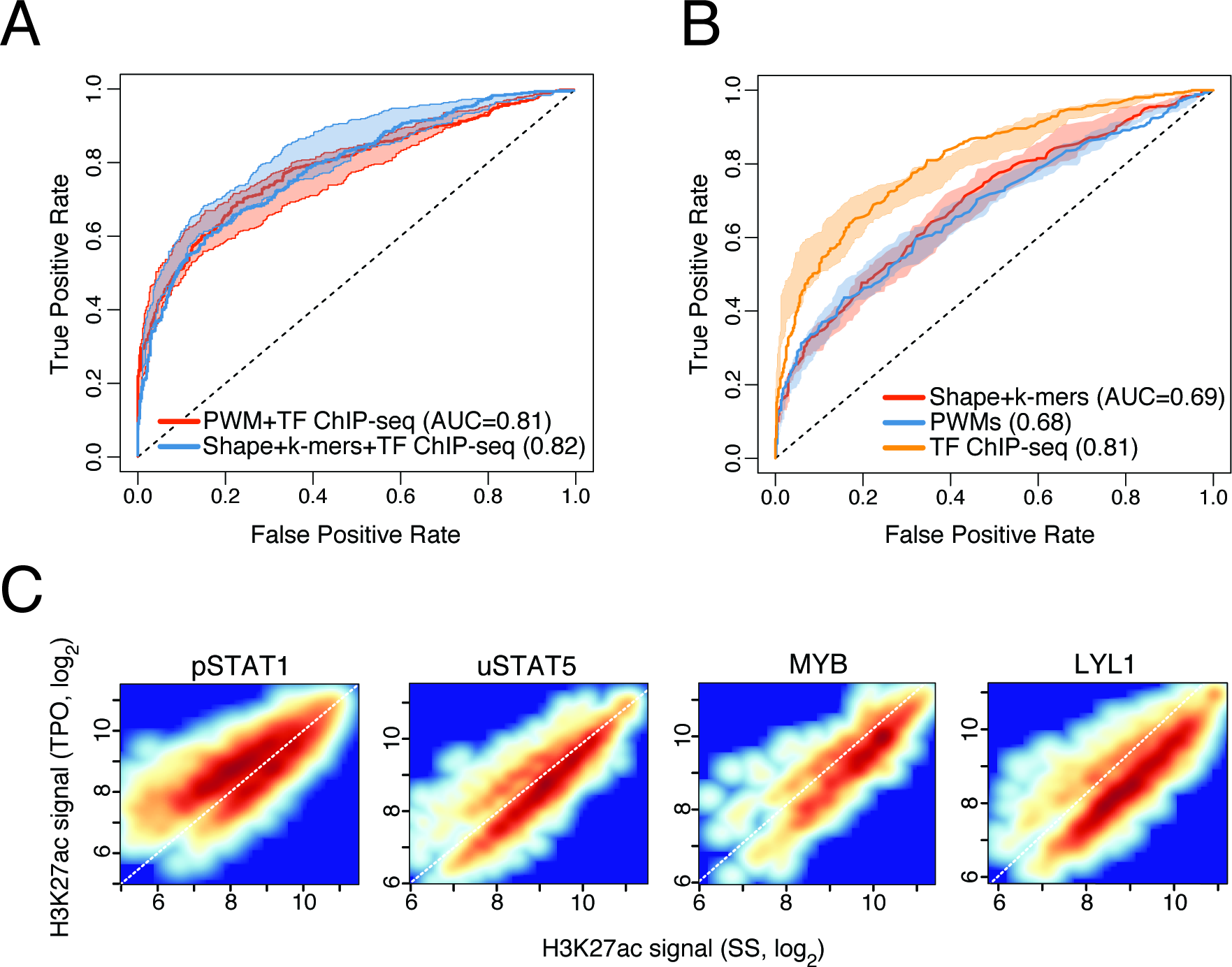
**(A)** Test set receiver operating characteristic (ROC) curve and area under the ROC curve (AUC) values for lasso models trained on the indicated sets of features. The shaded area is delimited by the ROC curves for models with highest and lowest AUC values, whereas the ROC curve for the model closest to mean AUC is shown. **(B)** Same as (A), for random forest classifiers. **(C)** Average normalized H3K27ac signals within ±4 kb of the summit of differentially acetylated DHSs bound by the indicated TFs. pSTAT1, tyrosine-phosphorylated STAT1; uSTAT5, tyrosine unphosphorylated STAT5. Supplemental Methods

## Supplemental Methods

### Subcellular RNA isolation

Subcellular fractionation and RNA preparation were performed essentially as described (Bhatt et al. 2012) with minor modifications. 1×10^6^ cells were serum starved and stimulated with TPO as above. Cells were collected by centrifugation and washed once with PBS. The cell pellet was resuspended in ice-cold NP-40 lysis buffer (10 mM Tris-HCl pH 7.5, 0.15% NP40, 150 mM NaCl). The lysate was then layered on 2.5 volumes of a sucrose buffer and centrifuged for 10 min at 13,000 rpm at 4°C. The supernatant (cytoplasmic fraction) was collected and added 3.5X volumes of RLT Buffer (Qiagen). The nuclei pellet was gently rinsed with ice-cold 1X PBS and resuspended in 200μl ice-cold glycerol buffer (20 mM Tris-HCl pH 7.9, 75 mM NaCl, 0.5 mM EDTA, 0.85 mM DTT, 0.125 mM PMSF, 50% glycerol) by gently flicking the tube. An equal volume of ice-cold nuclei lysis buffer (10 mM HEPES pH 7.6, 1 mM DTT, 7.5 mM MgCl2, 0.2 mM EDTA, 0.3 M NaCl, 1 M UREA, 1% NP-40) was added and gently vortexed twice for 2 sec, incubated for 2 min on ice, and then centrifuged at 13,000 rpm for 2 min at 4°C. The supernatant (nucleoplasmic fraction) was collected and added 3.5X volumes of RLT (Qiagen). The chromatin pellet was gently rinsed with cold 1X PBS and then dissolved in 500 μl TRIzol (Invitrogen). RNA purification from RLT-dissolved samples was performed using RNeasy columns (Qiagen). Chromatin-associated RNA was purified using Direct-zol (Zymo Research).

### Chromatin immunoprecipitation

Chromatin immunoprecipitation (ChIP) assays on serum-starved and TPO stimulated HPC-7 cells were performed as previously described (Wilson et al. 2010). Briefly, 1×10^8^ cells were cross-linked in 1% formaldehyde for 10 min at room temperature. Cells were lysed in 10mM Tris pH 8.0, 10mM NaCl and 0.2% NP40 containing inhibitors (1μg/mL leupeptin, 10mM NaBu and 50μg/mL PMSF) for 10 min on ice and were collected by centrifugation at 2500 rpm for 5 min at 4°C. The nuclei pellet was frozen until further use.

Frozen nuclei were resuspended in 50mM Tris pH 8.0, 10mM EDTA, 1% SDS supplemented with inhibitors (1μg/mL leupeptin, 10mM NaBu and 50μg/mL PMSF) and sonicated in an equal volume of IP dilution buffer (20mM Tris pH 8.0, 2mM EDTA, 150mM NaCl, 1% Triton X-100, 0.01% SDS and protease inhibitors) in ice-water (Bioruptor, Diagenode) for 5 cycles (30s on, 30s off). Chromatin was precleared with non-specific rabbit IgG (2 μg/μl, Sigma) for 1 hour and 100μL of protein G Dynabeads (Invitrogen) for 2 hours. The beads/IgG were removed by magnetic separation. Chromatin was immunoprecipitated at 4°C overnight using antibodies against H3K27ac (Abcam, 4729), histone H3 (Abcam, 1791), RAD21 (Abcam, 992), CTCF (Millipore, 07-729), STAT1 (Cell Signaling, 9172) or a control rabbit IgG (Invitrogen, 9172) and 100μl of protein G Dynabeads (Invitrogen) were added for additional 2 hours. Immunocomplexes were washed and eluted twice from the beads with 150μL elution buffer (100mM NaHCO3, 1% SDS). Cross-linking was reversed overnight with 0.3M NaCl and 2μL of RNase (10mg/mL) at 65°C, and samples were further treated with Proteinase K (20mg/mL) for 2 hours at 42°C. The ChIP DNA was purified using a PCR purification kit (Qiagen).

### Hi-C

Hi-C libraries were generated essentially as described (Schoenfelder et al. 2015) with modifications detailed below. 3.5×10^7^ HPC-7 cells were fixed in 2% formaldehyde (Agar Scientific) for 10 min, after which the reaction was quenched with ice-cold glycine (Sigma; 0.125M final concentration). Cells were collected by centrifugation (400 × g for 10 min at 4°C), and washed once with 50 mL PBS pH 7.4 (Gibco). After another centrifugation step (400 × g for 10 min at 4°C), the supernatant was completely removed and the cell pellets were immediately frozen in liquid nitrogen and stored at ‐80°C.

After thawing, the cell pellets were incubated in 50 mL ice-cold lysis buffer (10 mM Tris-HCl pH 8, 10 mM NaCl, 0.2% Igepal CA-630, protease inhibitor cocktail (Roche)) for 30 min on ice. After centrifugation to pellet the cell nuclei (650 × g for 5 min at 4°C), nuclei were washed once with 1.25 × NEBuffer 2 (NEB). The nuclei were then resuspended in 1.25 × NEBuffer 2, SDS (10% stock; Promega) was added (0.3% final concentration) and the nuclei were incubated at 37°C for one hour with agitation (950 rpm). Triton X-100 (Sigma) was added to a final concentration of 1.7 % and the nuclei were incubated at 37°C for one hour with agitation (950 rpm). Restriction digest was performed overnight at 37°C with agitation (950 rpm) with *HindIII* (NEB; 1500 units per 7 million cells). Using biotin-14-dATP (Life Technologies), dCTP, dGTP and dTTP (Life Technologies; all at a final concentration of 30 μM), the *HindIII* restriction sites were then filled in with Klenow (NEB) for 75 minutes at 37°C, followed by ligation for 4 hours at 16°C (50 units T4 DNA ligase (Life Technologies) per 7 million cells starting material) in a total volume of 5.5 mL ligation buffer (50 mM Tris-HCl, 10 mM MgCl_2_, 1 mM ATP, 10 mM DTT, 100 μg/mL BSA, 0.9 % Triton X-100) per 7 million cells starting material. After ligation, crosslinking was reversed by incubation with Proteinase K (Roche; 65 μl of 10mg/mL per 7 million cells starting material) at 65°C overnight. An additional Proteinase K incubation (65 μl of 10mg/mL per 7 million cells starting material) at 65°C for two hours was followed by RNase A (Roche; 15 μl of 10mg/mL per 7 million cells starting material) treatment and two sequential phenol/chloroform (Sigma) extractions. After DNA precipitation (sodium acetate 3M pH 5.2 (1/10 volume) and ethanol (2.5 × volumes)) overnight at ‐20°C, the DNA was spun down (centrifugation 3200 × g for 30 min at 4°C). The pellets were resuspended in 400 μl TLE (10 mM Tris-HCl pH 8.0; 0.1 mM EDTA), and transferred to 1.5 mL eppendorf tubes. After another phenol/chloroform (Sigma) extraction and DNA precipitation overnight at ‐20°C, the pellets were washed thrice with 70% ethanol, and the DNA concentration was determined using Quant-iT Pico Green (Life Technologies). For quality control, candidate 3C interactions were assayed (Supplemental Table S2) by PCR, and the efficiency of biotin incorporation was assayed by amplifying a 3C ligation product (Supplemental Table S2), followed by digest with *HindIII* or *NheI*.

To remove biotin from non-ligated fragment ends, 40 μg of Hi-C library DNA were incubated with T4 DNA polymerase (NEB) for 4 hours at 20°C, followed by phenol/chloroform purification and DNA precipitation overnight at ‐20°C. After one wash with 70% ethanol, sonication was carried out to generate DNA fragments with a size peak around 400 bp (Covaris E220 settings: duty factor: 10%; peak incident power: 140W; cycles per burst: 200; time: 55 seconds). After end repair (T4 DNA polymerase, T4 DNA polynucleotide kinase, Klenow (all NEB) in the presence of dNTPs in ligation buffer (NEB)) for 30 min at room temperature, the DNA was purified (Qiagen PCR purification kit). dATP was added with Klenow exo-(NEB) for 30 min at 37°C, after which the enzyme was heat-inactivated (20 min at 65°C). A double size selection using AMPure XP beads (Beckman Coulter) was performed: first, the ratio of AMPure XP beads solution volume to DNA sample volume was adjusted to 0.6:1. After incubation for 15 min at room temperature, the sample was transferred to a magnetic separator (DynaMag-2 magnet; Life Technologies), and the supernatant was transferred to a new eppendorf tube, while the beads were discarded. The ratio of AMPure XP beads solution volume to DNA sample volume was then adjusted to 0.9:1 final. After incubation for 15 min at room temperature, the sample was transferred to a magnet (DynaMag-2 magnet; Life Technologies). Following two washes with 70% ethanol, the DNA was eluted in 100 μl of TLE (10 mM Tris-HCl pH 8.0; 0.1 mM EDTA). Biotinylated ligation products were isolated using MyOne Streptavidin C1 Dynabeads (Life Technologies) on a DynaMag-2 magnet (Life Technologies) in binding buffer (5 mM Tris pH8, 0.5 mM EDTA, 1 M NaCl) for 30 min at room temperature. After two washes in binding buffer and one wash in ligation buffer (NEB), PE adapters (Illumina) were ligated onto Hi-C ligation products bound to streptavidin beads for 2 hours at room temperature (T4 DNA ligase NEB, in ligation buffer, slowly rotating). After washing twice with wash buffer (5 mM Tris, 0.5 mM EDTA, 1 M NaCl, 0.05% Tween-20) and then once with binding buffer, the DNA-bound beads were resuspended in a final volume of 90 μl NEBuffer 2. Bead-bound Hi-C DNA was amplified with 7 PCR amplification cycles using PE PCR 1.0 and PE PCR 2.0 primers (Illumina). After PCR amplification, the Hi-C libraries were purified with AMPure XP beads (Beckman Coulter). The concentration of the Hi-C libraries was determined by Bioanalyzer profiles (Agilent Technologies), and the Hi-C libraries were paired-end sequenced (HiSeq 4000, Illumina).

### Promoter Capture Hi-C

Promoter Capture Hi-C libraries were generated essentially as described (Schoenfelder et al. 2015) with modifications detailed below. 500 ng of Hi-C library DNA was resuspended in 3.6 μl H_2_O, and hybridization blockers (Agilent Technologies; hybridization blockers 1 and 2, and custom hybridization blocker) were added to the Hi-C DNA. Hybridization buffers and the custom-made RNA capture bait system (Agilent Technologies; designed as previously described (Schoenfelder et al. 2015): 39,021 individual biotinylated RNAs targeting the ends of 22,225 promoter-containing mouse *HindIII* restriction fragments) were prepared according to the manufacturer’s instructions (SureSelect Target Enrichment, Agilent Technologies). The Hi-C library DNA was denatured for 5 min at 95°C, and then incubated with hybridization buffer and the RNA capture bait system at 65°C for 24 hours (all incubation steps in a MJ Research PTC-200 PCR machine). After hybridization, 60 μl of MyOne Streptavidin T1 Dynabeads (Life Technologies) were washed thrice with 200 μl binding buffer (SureSelect Target Enrichment, Agilent Technologies), before incubation with the Hi-C DNA/RNA capture bait mixture with 200 μl binding buffer for 30 min at room temperature, slowly rotating. Hi-C DNA bound to capture RNA was isolated using a DynaMag-2 magnet (Life Technologies). Washes (15 min in 500 μl wash buffer I at room temperature, followed by three 10 min incubations in 500 μl wash buffer II at 65°C) were performed according to the SureSelect Target enrichment protocol (Agilent Technologies). After the final wash, the beads were resuspended in 300 μl NEBuffer 2, isolated on a DynaMag-2 magnet, and then resuspended in a final volume of 30 μl NEBuffer 2. After a post-capture PCR (four amplification cycles using Illumina PE PCR 1.0 and PE PCR 2.0 primers; 13 to 15 individual PCR reactions), the Promoter CHi-C libraries were purified with AMPure XP beads (Beckman Coulter). The concentration of the Promoter CHi-C libraries was determined by Bioanalyzer profiles (Agilent Technologies), and the Promoter CHi-C libraries were paired-end sequenced (HiSeq 4000, Illumina).

### RNA-seq data analysis

Subcellular RNA-seq reads were quality and adapter trimmed using Trimmomatic v0.33 (Bolger et al. 2014) in palindrome mode (Illumina TruSeq universal adapter sequence). Reads were aligned to the mouse reference genome (GRCm38 primary assembly, release 82) using STAR v2.4.2a (Dobin et al. 2013). Gene-level count tables were generated while mapping based on Gencode vM7 annotations. Normalized (reads per million, RPM) strand-specific bedGraph tracks were generated while mapping and converted to bigWig format using BEDTools v2.17.0 (Quinlan and Hall 2010). For visualization, the RNA-seq coverage was log transformed with a pseudocount of 1 where indicated.

Downstream analyses were performed using R v3.3.2 and Bioconductor (Huber et al. 2015). Differentially transcribed and differentially expressed genes were identified using DESeq2 v1.14.1 (Love et al. 2014). False discovery rates were controlled at a

0. 1% level by applying an independent hypothesis weighting (IHW) procedure (Ignatiadis et al. 2016) using the mean of normalized counts for each gene as the informative covariate. Conditional independence of p-values under the null hypothesis was verified prior to IHW.

### ChIP-seq data analysis

ChIP-seq reads were aligned to the mouse reference genome (GRCm38 primary assembly, release 82) using Bowtie2 v2.2.3 (Langmead and Salzberg 2012), allowing no mismatch within a seed of 22 nt. SAM files were converted to BAM format, sorted, de-duplicated and indexed with Sambamba v0.6.5 (Tarasov et al. 2015). Reads mapping to unplaced scaffolds or to the mitochondrial genome were discarded. Downstream analyses were performed using Homer v4.9 (Heinz et al. 2010). Uniquely aligned reads were used to generate Homer tag directories, and peak calling was performed using the findPeaks function. For H3K27ac ChIP-seq profiles, peaks were called against matched IgG control profiles in histone mode with a 1 kb window size and a 2.5 kb distance cutoff. High-confidence peak regions were defined as the peak intersection between biological replicates (computed with BEDTools v2.17.0, minimum overlap of 1 bp). Normalized (RPM) bigWig tracks were generated using the makeMultiWigHub.pl script.

### Classification of cis-regulatory elements

DNaseI-seq data (Wilson et al. 2016) were aligned to the mm10 mouse reference genome with Bowtie2 v2.2.3, allowing no mismatch within a 22 nt seed. DNaseI hypersensitive sites (DHS) were called from a pool of four biological replicates using MACS2 v2.1.0 (Zhang et al. 2008) and ‐‐no-model option, at 5% FDR. DHS separated by less than 150 bp (approximately one nucleosome) were merged, and H3K4me1 and H3K4me3 signals, defined as offset-normalized enrichments over input, were computed in 1 kb windows centered on the resulting DHS regions. This window size maximized the variance of H3K4me1/H3K4me3 ratios among all tested window sizes in the range (10, 2500) bp. DHS exhibiting an H3K4me1 and H3K4me3 fold enrichment over input < 2 were classified as insulators and were not considered further. Offset-normalized H3K4me1/H3K4me3 ratios × were computed at the remaining DHS regions. To discriminate enhancers from promoters, the distribution of log2(X) was fitted by a two-component Gaussian mixture model with unequal variance using mclust v5.1.

### Differential acetylation analysis

Differentially acetylated regions (DARs) were called from the union of all high-confidence H3K27ac peaks identified across conditions using csaw v1.8.0 (Lun and Smyth 2015) with the following parameters: window size, 1000 bp; spacing, 200 bp; fragment length, 200 bp; filter, 10 reads; minimum mapping quality, 10. Differentially acetylated windows were identified by fitting a quasi-likelihood negative binomial model (Lun et al. 2016) and subsequently merged into DARs at 1% FDR. DARs exhibiting heterotypic differentially acetylated windows were discarded. DARs were annotated to genomic compartments and cis-regulatory elements using the ChIPseeker package v1.10.3 (Yu et al. 2015).

### Hi-C data analysis

Hi-C reads were pre-processed with HiC-Pro v2.7.7 (Servant et al. 2015). The digest_genome.py utility was used to digest the mouse reference genome (GRCm38 primary assembly, release 82) in silico, generating a BED file containing all theoretical HindIII restriction fragments. HiC-Pro was run with Bowtie2 v2.2.3. The algorithm performs a two-step read alignment to the reference genome whereby reads are first independently aligned using the Bowtie2 end-to-end algorithm, followed by detection of ligation sites on unmapped chimeric reads and re-alignment of their 5' fractions. Singletons, multi-mappers, duplicated reads, and read pairs with mapping quality < 10 were discarded. Aligned reads were assigned to restriction fragments and read pairs corresponding to invalid ligation products (e.g. self-ligations, dangling ends) or mapping outside the insert size range [150, 800], or anchored at unplaced scaffolds or the mitochondrial genome were discarded. After a correlation analysis of Hi-C signals between individual samples, valid pairs from biological replicates were merged. These were used to compute raw and coverage-and-distance corrected ICE (iterative correction and eigenvector decomposition) contact matrices (Imakaev et al. 2012), and to generate Homer tag directories for which signals exceeding 10 times the average read counts in 10 kb bins were removed.

A/B compartments and topological domains were called using Homer as follows. For each chromosome, A/B compartments were identified by computing the first eigenvector of a binned (40kb resolution) interaction profile correlation matrix. Transcription start site coordinates of annotated genes were used to assign active and inactive compartments to positive and negative eigenvector values, respectively. TADs were identified using the Homer Hi-C domain finding algorithm, which computes the ratio of upstream and downstream interaction counts (directionality index, DI) within a given fixed-size window (Dixon et al. 2012). A window size of 1 Mb, a bin size of 25 kb and a step size of 5 kb were used. Bins exhibiting coverage values smaller than 15% of the mean bin coverage or exceeding it by more than four standard deviations were excluded. DI values were smoothed using a running average over ±25 kb window. Domain boundary coordinates were defined based on smoothed DI profiles, requiring a minimum index score of 0.5. For comparison of TAD coordinates across conditions, a maximum tolerance of 10 kb was allowed at the TAD boundaries. Differentially acetylated TADs were identified using csaw as described above. Differentially acetylated windows were merged into differentially acetylated TADs at 1% FDR, using the genomic coordinates of TADs identified in basal condition and requiring at least 75% homotypic differentially acetylated windows therein.

To identify CTCF loops, statistically significant Hi-C interactions were first called using the Homer analyzeHiC algorithm with parameters-interactions ‐res 10000 ‐superRes 20000 ‐center ‐nomatrix. Significant interactions were then filtered to retain only intrachromosomal interactions at 5% FDR. Of these, only interactions whose anchors overlapped both a CTCF and a Rad21 peak and exhibited an unambiguous CTCF motif direction were considered. CTCF loops were then defined as CTCF/Rad21-anchored interactions with convergent CTCF motif orientation.

### Structured interaction matrix analysis

Enhancer-enhancer interactions within and between differentially acetylated TADs were analyzed using a structured interaction matrix analysis (SIMA) (Lin et al. 2012), which pools Hi-C signals across genomic regions of interest located within a given set of chromatin domains. SIMA was run separately on activated and repressed TADs of size ≥ 200 kb (-minDsize 2e5) to analyze two sets of genomics regions of interest: i) differentially acetylated enhancers (i.e. enhancers located within DARs, resized to 1 kb); ii) a control set of genomic regions obtained by systematically shifting (10 kb downstream) the genomic coordinates of differentially acetylated enhancers. A superresolution of 10 kb (-superRes 10000; i.e. all reads within a 10 kb window centered on each enhancer were considered) and a resolution of 2.5 kb (-res 2500) were used. Interactions across TADs were analyzed for differentially acetylated TADs separated by ≤ 20 Mb (-max 2e7). The same analysis was performed for CTCF loops.

### Promoter Capture Hi-C data analysis

Promoter Capture Hi-C (PCHi-C) reads were pre-processed with HiC-Pro v2.7.7 as described above. Statistically significant PCHi-C interactions were computed with CHiCAGO v1.2.0 (Cairns et al. 2016). To this end, a restriction map file and a bait map file (both in BED format) were used to precompute auxiliary files using the makeDesignFiles.py script from chicagoTools. To generate input files for the CHiCAGO pipeline, HiC-Pro valid pairs in BAM format were mapped to restriction fragments using the HiC-Pro mapped_2hic_fragments.py script, de-duplicated using Sambamba v0.6.5, sorted by natural sort, and converted to CHiCAGO format using the bam2chicago.sh script provided by chicagoTools. CHiCAGO design files were generated using the makeDesignFiles.py script with parameters ‐‐minFragLen=150 ‐‐ maxFragLen=40000 ‐‐maxLBrownEst=1500000 ‐‐binsize=20000 ‐‐removeb2b=True ‐‐removeAdjacent=True. An interaction score ≥ 5 was used to call statistically significant PCHi-C interactions. Annotation of PCHi-C interactions to genomic regions was performed using the GenomicInteractions package v1.8.0 (Harmston et al. 2015). To correlate changes in H3K27ac levels with changes in PCHi-C interaction frequency at individual differentially transcribed genes, cis-regulatory units were first defined for each baited promoter by considering the set of all its PIRs across conditions. PIRs were further filtered to retain only regions overlapping at least one high-confidence H3K27ac peak, ensuring stable estimates of H3K27ac ratios between conditions. In addition, cis-regulatory units containing less than five H3K27ac-marked PIRs were discarded, resulting in a set of 907 differentially transcribed loci that were further analyzed as follows. Given a cis-regulatory unit, the normalized H3K27ac fold change and the normalized ratio between PCHi-C signals in TPO and basal conditions (interaction frequency fold change) were computed for each PIR. The correlation between interaction frequency and H3K27ac fold changes was then computed for each cis-regulatory unit using a Spearman’s rank correlation coefficient. Statistically significant correlation values were identified by deriving a null distribution for the correlation coefficient. This was estimated using a randomization procedure whereby the observed (H3K27ac fold change, interaction frequency fold change) pairs were randomly permuted 50 times for each cis-regulatory unit and the corresponding Spearman’s rank correlation coefficient were recorded. The empirical distribution of all recorded correlation values was used to perform a one-sided statistical test.

### Superenhancers

Superenhancers were defined starting from the genomic coordinates of the individual constituent enhancer elements. First, enhancers located within 2.5 kb of annotated transcription start sites or not overlapping an H3K27ac peak were discarded. The remaining enhancers were then stitched together if located closer than 20 kb. This value has been chosen after examining the monotone relation between the number of stitched regions and the distance threshold. Stitched regions were then ranked by total input-normalized H3K27ac signals and classified in superenhancers or regular enhancers as previously described (Whyte et al. 2013).

Differentially acetylated SEs were identified using csaw as described above. Differentially acetylated windows were merged into differentially acetylated SEs at 1% FDR, using the genomic coordinates of SEs identified in basal condition. Differentially acetylated SEs exhibiting heterotypic differentially acetylated windows were discarded.

### Motif analysis

De novo motif discovery was performed using Homer (findMotifsGenome.pl script), searching for motifs of length 6-12 nt. For the analysis of differentially acetylated cis-regulatory elements, DHSs located within DARs were considered and 200 nt sequences centered on the DHS summit were extracted. For motif density analysis, PWMs of top TF motif hits and a collection of 363 Homer PWMs for vertebrate TFs were scored within 2kb windows centered on DHS summits at 10 bp resolution using the annotatePeaks.pl script.

### Statistical learning

Logistic regression models are a class of probabilistic binary classifiers. Least absolute shrinkage and selection operator (lasso) logistic regression penalizes model complexity through an L1 norm penalty. The lasso generates a sparse model representation through intrinsic feature selection, effectively contrasting overfitting in high-dimensional feature spaces (Tibshirani 1996). Model parameters are usually estimated from a training set of labeled data points using cross-validation, and model performances are evaluated on a test set that has not been previously seen by the model.

Here, lasso logistic regression models were used to predict rapid cis-regulatory responses to TPO signaling. For feature scoring, DNA sequences of differentially acetylated cis-regulatory elements (defined as promoters and enhancers located within DARs, see above) were extracted from the mm10 reference genome within 200 bp and 500 bp windows centered on the DHS summit. The following features were scored therein:

1. DNA sequence content encoded as *k*-mers (2 ≤ *k* ≤ 4), computed within 500 bp windows.
2. Average DNA shape feature values within 200 bp windows. These features were computed using the R package DNAshapeR v1.2.0 (Chiu et al. 2016), which implements a high-throughput approach based on all-atom Monte-Carlo predictions. Four DNA shape features were used in this study: helix twist, propeller twist, minor groove width and roll.
3. Transformed FIMO p-values (-10*log_10_(p)) (Grant et al. 2011) for a curated collection of >1,700 single and composite position weight matrices (PWMs) representing mammalian TF motifs (Diaferia et al. 2016). Only TFs expressed in HPC-7 cells were considered (FPKM ≥ 1 across all cytoplasmic RNA-seq samples), along with composite motifs for expressed TFs. FIMO scores were computed as previously described (Barozzi et al. 2014) using the FIMO implementation provided as part of MEME v4.11.3.
4. ChIP-seq signals for a collection of 29 genome-wide binding profiles for hematopoietic and other sequence-specific TFs in HPC-7 cells (Supplemental Table S1). These profiles were generated as part of this study or previously published (Wilson et al. 2010, 2016). Feature enrichments were computed as previously described (Comoglio et al. 2015) within 1 kb windows centered on the DHS summit.
5. Normalized Hi-C signals defined as the total number of valid di-tags anchored within 1 kb windows centered on the DHS summit. These were used as a proxy for interaction frequencies.

Data points were randomly partitioned into 100 balanced training (80%) and test (20%) sets composed of an equal number of activated and repressed cis-regulatory elements. Lasso logistic regression models were trained on each training set with tenfold cross validation using the glmnet implementation (Friedman et al. 2010). The value of the regularization parameter that minimized the cross-validated misclassification error was used to predict the class labels of the cis-regulatory elements in the corresponding test set. Model performances were evaluated by computing the area under the receiver operating characteristic curve (AUC) using the R package ROCR v1.0-7 (Sing et al. 2005) and average AUC values across all 100 models were computed. Feature importance analysis was carried using the bootstrap-lasso algorithm as previously described (Comoglio and Paro 2014; Comoglio et al. 2015). Features with selection probability (stability) ≥ 0.7 were considered.

Random forest classifiers (Breiman 2001) were trained and evaluated on the same balanced training and test sets using the R package randomForest v4.6-12.

